# BrainBeats, an Open-Source EEGLAB Plugin to Jointly Analyze EEG and Cardiovascular Signals

**DOI:** 10.1101/2023.06.01.543272

**Authors:** Cédric Cannard, Helané Wahbeh, Arnaud Delorme

## Abstract

The interplay between the brain and the cardiovascular systems is garnering increased attention for its potential to advance our understanding of human physiology and improve health outcomes. However, the multimodal analysis of these signals is challenging due to the lack of guidelines, standardized signal processing and statistical tools, graphical user interfaces (GUIs), and automation for processing large datasets and increasing reproducibility. A further void exists in standardized quantitative EEG (qEEG) and heart-rate variability (HRV) feature extraction methods, undermining clinical diagnostics or the robustness of machine learning (ML) models. In response to these limitations, we introduce the BrainBeats toolbox. Implemented as an open-source EEGLAB plugin, BrainBeats integrates three main protocols: 1) Heartbeat-evoked potentials (HEP) and oscillations (HEO) for assessing time-locked brain-heart interplay at the millisecond accuracy; 2) qEEG and HRV feature extraction for examining associations/differences between various brain and heart metrics or for building robust feature-based ML models; 3) Automated extraction of heart artifacts from EEG signals to remove any potential cardiovascular contamination while conducting EEG analysis. We provide a step-by-step tutorial for performing these three methods on an open-source dataset containing simultaneous 64-channel EEG, ECG, and PPG signals. Users can easily fine-tune parameters to tailor their unique research needs via the graphical user interface (GUI) or the command line. BrainBeats should make brain-heart interplay research more accessible and reproducible.

**SUMMARY:** The BrainBeats toolbox is an open-source EEGLAB plugin designed to jointly analyze EEG and cardiovascular (ECG/PPG) signals. It offers three main protocols: heartbeat-evoked potentials (HEP) assessment, feature-based analysis, and heart artifact extraction from EEG signals. It should aid researchers and clinicians in studying brain-heart interplay through two lenses (HEP and features), enhancing reproducibility and accessibility.

## INTRODUCTION

For a long time, the reductionist approach has dominated scientific inquiry in human physiology and cognition. This approach involved dissecting complex bodily and mental processes into smaller, more manageable components, allowing researchers to focus on individual systems in isolation. This strategy arose due to the immense challenge of studying the intricate and interconnected nature of the human body and mind^1^. Reductionism has been instrumental in understanding individual subsystems in isolation, such as elucidating the role of ion channels and action potentials for neural^2^ or cardiac^3^ communication. However, a significant gap remains in understanding how these isolated systems interact on a larger spatial and temporal scale. The multimodal (integrative or ecological) framework considers the human body a complex multidimensional system, where the mind is seen not as a product of the brain but as an activity of the living being, an activity that integrates the brain within the everyday functions of the human body^4^. The multimodal and reductionist approaches are not exclusive, just like we cannot study one neuron without the whole brain or the whole brain without understanding individual neuron properties. Together, they pave the way for a more comprehensive, synergetic understanding of human health, pathology, cognition, psychology, and consciousness. The present method aims to ease the multimodal investigation of the interplay between the brain and the heart by providing joint analysis of electroencephalography (EEG) and cardiovascular signals, namely electrocardiography (ECG) and photoplethysmography (PPG). This toolbox, implemented as an EEGLAB plugin in MATLAB, addresses existing methodological limitations and is made open source to facilitate accessibility and reproducibility in the scientific area. It implements the latest guidelines and recommendations into its design and default parameters to encourage users to follow known best practices. The proposed toolbox should be a valuable resource for researchers and clinicians interested in 1) extracting features from EEG and ECG/PPG signals, 2) studying heartbeat-evoked potentials, or 3) removing heart artifacts from EEG signals.

### Heart-brain research: which measures?

The relationship between the heart and the brain has been historically studied via neuroimaging methods such as functional magnetic resonance imaging (fMRI) and positron emission tomography (PET). Using these tools, researchers highlighted some brain regions associated with cardiovascular control (e.g., manipulation of heart rate and blood pressure^5^), showed the influence of heart rate on the BOLD signal^6^, or identified potential brain-body pathways contributing to coronary heart disease (i.e., stress-evoked blood pressure^7^). While these studies have significantly advanced our understanding of the complex interplay between the central nervous system (CNS) and cardiovascular function, these neuroimaging techniques are expensive, have limited availability, and are confined to controlled laboratory settings, which restricts their practicality for real-world and large-scale applications.

In contrast, EEG and ECG/PPG are more affordable and portable tools that offer the potential for studying brain-heart interactions in more diverse settings and populations or over longer periods, providing new opportunities. ECG measures the electrical signals generated by each heartbeat when the heart contracts and relaxes via electrodes placed on the skin (usually on the chest or arms)^8^. PPG measures blood volume changes in the microvascular tissues (i.e., blood flow and pulse rate) using a light source (e.g., LED) and a photodetector (commonly placed on a fingertip, wrist, or forehead), relying on how blood absorbs more light than the surrounding tissue^9^. Both methods provide valuable information about cardiovascular function but serve different purposes and offer distinct data types. Like ECG, EEG records the electrical fields generated by the synchronized activity of thousands of cortical neurons that propagate through the extracellular matrix, tissues, skull, and scalp until they reach the electrodes placed on the scalp’s surface^10^. As such, the use of EEG and ECG/PPG holds great promise for advancing our understanding of the physiological, cognitive, and emotional processes underlying brain-heart interactions and their implications for human health and well-being. Therefore, capturing heart-brain interplay from EEG, ECG/PPG signals with the BrainBeats toolbox may be particularly useful for the following scientific areas: clinical diagnostic and forecasting, big data machine learning (ML), real-world self-monitoring^11^, and mobile brain/body imaging (MoBI)^12,13^.

### Two approaches for jointly analyzing these signals

There are two main approaches to studying interactions between EEG and cardiovascular signals:

1. The heartbeat-evoked potentials (HEP) in the time domain: event-related potentials (ERP). And the heartbeat-evoked oscillations (HEO) in the time-frequency domain: event-related spectral perturbations (ERSP) and inter-trial coherence (ITC). This approach examines how the brain processes each heartbeat. With millisecond (ms) accuracy, this method requires that both time series are perfectly synchronized and that the heartbeats are marked in the EEG signals. This approach has gained interest in recent years^14–19^.
2. Feature-based: this approach extracts EEG and heart-rate variability (HRV) features from continuous signals and examines associations between them. This has been done independently for EEG (often termed quantitative EEG or qEEG^20^), ECG^21–23^, and PPG^24–26^. This approach presents promising applications by capturing both state-and trait-related variables. Note that, for both EEG and cardiovascular signals, the longer the recording, the more dominant the trait variable^27–29^. Thus, the applications depend on the recording parameters. Feature-based analyses are gaining growing interest, providing new quantitative metrics for forecasting the development of mental and neurological disorders, treatment-response, or relapse^30–35^. This approach is especially compelling with large and real-world datasets (e.g., clinic, remote monitoring), which can be more easily obtained thanks to the recent innovations in wearable neurotechnology^11^. A less explored application is the identification of associations between specific brain and heart features, highlighting potential underlying central nervous system dynamics. Heart rate variability (HRV) can be calculated from both ECG and PPG signals. It provides information about the autonomous nervous system (ANS) by measuring the variations in time intervals between heartbeats (i.e., the normal-to-normal intervals)^27^. Increased sympathetic (SNS) activity (e.g., during stress or exercise) typically reduces HRV, while parasympathetic (PNS) activity (e.g., during relaxation) increases it. A slower breathing rate generally increases HRV due to enhanced PNS activity, especially for short recordings (<10 min)^27^. Higher HRV scores generally suggest a more resilient and adaptable ANS, while a lower HRV can indicate stress, fatigue, or underlying health issues. Long HRV recordings (i.e., at least 24 h) provide a predictive prognosis for various health conditions, including cardiovascular diseases, stress, anxiety, and some neurological conditions^27^. Measures like blood pressure, heart rate, or cholesterol levels give information about the cardiovascular system’s status. In contrast, HRV adds a dynamic aspect, showing how the heart responds to and recovers from stress.

### BrainBeats’ advantages over existing methods

While many tools exist to process cardiovascular and EEG signals independently from one another, none is currently available for jointly analyzing them. Furthermore, most available means to process cardiovascular signals involve costly licensing, do not allow automated processing (especially beneficial for large datasets), have proprietary algorithms that prevent transparency and reproducibility, or require advanced programming skills by not providing a graphical user interface (GUI)^36^. To our knowledge, four open-source MATLAB toolboxes support HEP/HEO analysis with a GUI: the ecg-kit toolbox^37^, the BeMoBIL pipeline^38^, the HEPLAB EEGLAB plugin^39^, and the CARE-rCortex toolbox^40^. While HEPLAB, BeMoBIL, and ecg-kit facilitate HEP analysis by detecting heartbeats and marking them in the EEG signals, they do not provide statistical analysis or are limited to the time domain (i.e., HEP). The CARE-rCortex plugin addressed these issues by supporting ECG and respiratory signals, time-frequency domain analysis, statistics, and advanced baseline normalization and correction methods adapted to HEP/HEO analysis. However, it uses the Bonferroni method for statistical correction of the type 1 error (i.e., false positives), which is too conservative and not physiologically sound for EEG applications, leading to an increase in type II errors (i.e., false negatives)^41^. Furthermore, the toolbox does not offer command-line access for automation. Finally, recent studies recommend against baseline correction methods^42–44^, as they reduce the signal-to-noise ratio (SNR) and are statistically unnecessary and undesirable.

To address these limitations, we introduce the BrainBeats toolbox, currently implemented as an open-source EEGLAB plugin in the MATLAB environment. It incorporates the following advantages over previous methods:

1. An easy-to-use GUI and command-line capabilities (for programmers aiming to perform automated processing).
2. Validated algorithms, parameters, and guidelines for processing cardiovascular signals, such as detecting R peaks, interpolating RR artifacts, and computing HRV metrics (e.g., implanting guidelines for windowing, resampling, normalization, etc.^27,45,46^). This is important because Vest et al. demonstrated how modest differences in these processing steps can lead to divergent results, contributing to the lack of reproducibility and clinical applicability of HRV metrics^46^.
3. Validated algorithms, default parameters, and guidelines for processing EEG signals, including filtering and windowing^44,47^, re-referencing^48,49^, removal of abnormal channels and artifacts^50–52^, optimized ICA decomposition and classification of independent components^53–56^. The users can fine-tune all preprocessing parameters or even preprocess their EEG data with their preferred method before using the toolbox to match their needs (e.g., with EEGLAB clean_rawdata plugin^50,52^, the BeMoBIL pipeline^38^, the PREP pipeline^57^, etc.).
4. Heartbeat-evoked potentials (HEP, i.e., time domain) and oscillations (HEO; event-related spectral perturbations with wavelet or FFT methods, and inter-trial coherence are available through the standard EEGLAB software) from ECG signals. Parametric and nonparametric statistics with corrections for type 1 errors are available via EEGLAB’s standard software. Nonparametric statistics include permutation statistics and spatiotemporal corrections for multiple comparisons (e.g., spatiotemporal clustering or threshold-free cluster enhancement)^58,59^. Users can use the LIMO-EEG plugin to implement hierarchical linear modeling, which accounts well for within and between-subjects variance and implements an assumption-free mass-univariate approach with robust control for type I and II errors^60,61^. The HEP/HEO data statistical analyses can be performed in the channel and independent component domains.
5. HEP/HEO and HRV analysis from PPG signals (for the first time for HEP/HEO).
6. Supports the joint extraction of EEG and HRV features for the first time.
7. The toolbox provides various data visualizations to inspect signals at various necessary processing steps and outputs at the subject level.

**TABLE 1.** Novelties brought by BrainBeats relative to pre-existing, similar methods.

| METHOD | Detect R-peaks from ECG | Detect R-waves from PPG | HEP/HEO | EEG & HRV features | Remove heart artifacts from EEG | GUI | Command line |
| --- | --- | --- | --- | --- | --- | --- | --- |
| ecg-kit | X | X | X |  |  |  | X |
| BeMoBIL | X |  | X |  |  |  | X |
| HEPLAB | X |  | X |  |  | X | X |
| CARE-rCortex | X |  | X |  |  | X | X |
| BrainBeats | X | X | X | X | X | X | X |

### Information to help readers decide whether the method is appropriate for them

This toolbox is appropriate for any researcher or clinician with EEG and ECG/PPG data. The plugin does not yet support importing EEG and ECG/PPG signals from separate files (although this feature will be available soon). The toolbox is appropriate for anyone aiming to perform HEP/HEO analysis, extract EEG and/or HRV features with standardized methods, or simply remove heart artifacts from EEG signals. See Diagram for a summary of BrainBeats overall flow and methods.

**DIAGRAM.**
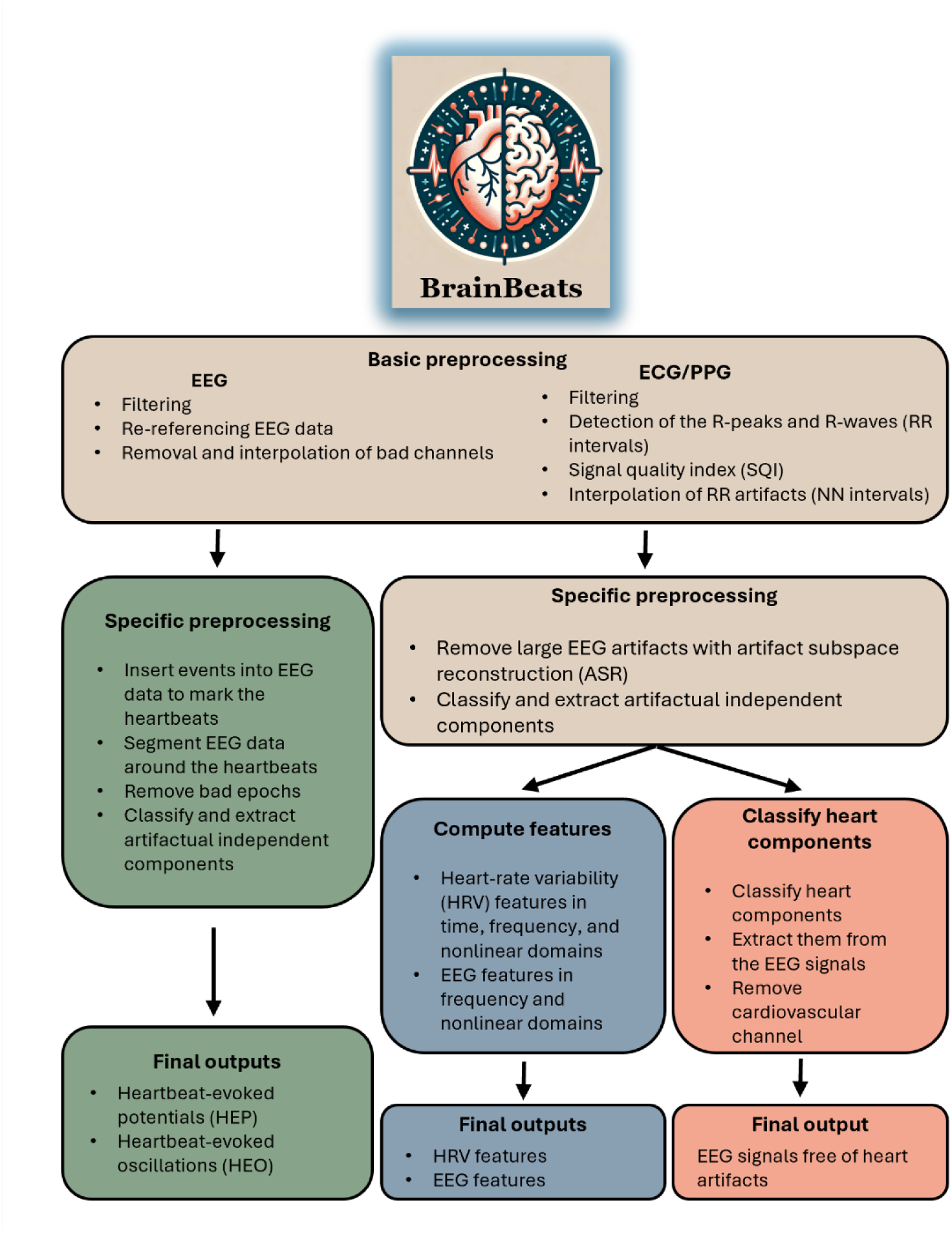
Block diagram summarizing BrainBeats’ overall architecture and flow. The operations that are common across the three methods are brown. Operations specific to heartbeat-evoked potentials (HEP) and oscillations (HEO) are green. Operations specific to the extraction of EEG and HRV features are blue. Operations specific to removing heart artifacts from the EEG signals are red.

#### PROTOCOL

**1. BrainBeats requirements**
  1.1. Install MATLAB and EEGLAB on the computer. EEGLAB can be downloaded at https://github.com/sccn/eeglab and unzipped (or cloned for Git users) anywhere on the computer. See the GitHub page for more details on installation.
  1.2. Add the path to the EEGLAB folder in MATLAB’s home panel by clicking **Set Path** button. Select the eeglab folder with the unzipped file and click **Save > Close**.
  1.3. Launch EEGLAB by typing **eeglab** in MATLAB’s command window.
  1.4. Install the BrainBeats plugin by clicking **File > Manage EEGLAB extensions**. Type
  **brainbeats** in the search bar, select the **BrainBeats** plugin in the list and click **Install/Update**.
  1.5. Load the sample dataset into EEGLAB. Click **File > Load existing data**. Navigate to the EEGLAB folder, go to the plugins folder, go to the BrainBeats folder and open to the sample_data folder. Select the file **dataset.set**. NOTE: This dataset corresponds to sub-032 (resting state with eyes opened) from the open-source dataset^62,63^. Informed consent was obtained from each participant, and the Ural Federal University ethics committee approved the experimental protocol. We selected this because it contains simultaneous EEG (64 channels), ECG (one channel), and PPG (one channel), acquired using an ActiCHamp amplifier (Brain Products, Germany). ECG and PPG signals were collected using the auxiliary inputs. These data were collected at a sampling rate of 1000 Hz, and no online digital filtering. EEG data were recorded with active electrodes placed according to the 10–20 system, with FCz as the online reference and Fpz as the ground electrode, and with the impedance maintained below 25 kOm. ECG was recorded using one active electrode placed on the right wrist, the reference electrode on the left wrist, and the ground on the left inner forearm at 3 cm distally from the elbow. PPG was recorded from the left index finger. EEG, ECG, and PPG data are time-synchronized since they were recorded simultaneously with the same amplifier. See references^62,63^ for more details. For this tutorial, we merged the EEG, ECG, and PPG data into one EEGLAB dataset, loaded the 3D boundary element method (BEM) electrode coordinates, and downsampled the signal to 250 Hz to reduce the file weight (for user download) and accelerate computing time. Because this dataset had no abnormal EEG channel, we artificially modified channel TP9 to illustrate BrainBeats’ bad channel detection and removal algorithm. Similarly, we artificially simulated a large EEG artifact at the beginning of the file and a high-frequency muscle-like artifact in the temporal channels from 3–6 s to illustrate BrainBeats’ artifact removal feature.
  1.6. Check the **Save outputs** checkbox to save everything in the corresponding .set file at the same location as the original file that was loaded into EEGLAB.
**2. Heartbeat-evoked potentials (HEP) and oscillations (HEO)**
  2.1. Open BrainBeats’ first general user interface (GUI) to select the main parameters. In EEGLAB, click **Tools > BrainBeats > 1^st^ level (subject level).** Select **Heartbeat-evoked potentials HEP** as the analysis to run, **ECG** as the heart data type, and click on the button to display the list of channels to select the ECG channel labeled ECG (or type it directly in the text box). Keep the options **Visualize outputs (recommended)** and **Save outputs** selected and click **Ok**. See **Figure 1**.
  2.2. A second GUI window pops up based on the previous choices (i.e., HEP analysis with ECG data). Select preprocessing parameters for both EEG and ECG signals. You can turn off preprocessing by unchecking the boxes **Preprocess ECG** and **Preprocess EEG** if your data were already preprocessed prior to launching BrainBeats (**Figure 2**). Change **Power line noise** to **50 Hz** in the **Preprocess EEG section** since these EEG data were recorded in Russia. Click **Ok** to launch.
  2.3. A warning message will appear, asking for confirmation to remove the PPG channel that has been detected (**Figure 3**). This is because the toolbox is not designed to analyze both ECG and PPG simultaneously (or other auxiliary channels), and keeping it in the dataset will lead to severe errors (e.g., artifact removal, poor ICA decomposition, etc.). Click **Yes**. BrainBeats starts performing some checks, setting some default parameters, and separates the ECG from the EEG data to preprocess the ECG signal and compute the RR intervals.
  2.4. ECG and RR time series are preprocessed using validated algorithms from the Physionet Cardiovascular Signal toolbox^46,64^ adapted to fit BrainBeats’ data formatting, increase clarity, parameter tuning, and computing time (see references for validation of the algorithms). The plugin outputs the RR intervals, timestamps, filtered ECG signal, R-peaks indices, and heart rate (HR). Tune these parameters via the GUI (**Figure 2**) or command line. NOTE: The ECG signal is bandpass-filtered using a customized, validated filter (1-30 Hz) and scans the signal to identify the QRS complex and R-peaks using the Pan–Tompkins (P&T) method^65^, implementing some signal processing operations including differentiation, squaring, integration, and smoothing for best performance. The P&T energy threshold is estimated based on the sample rate and the smoothed ECG values to avoid disruption from large bumps. If the RR interval variability exceeds 1.5 times the median, it searches for missed peaks. The mean R-peak sign is calculated over 30 s segments, and peak points are refined through a refractory period check, managing flatline conditions and ensuring consistent detection.
  2.5. Next, BrainBeats identifies abnormal RR intervals or spikes within RR intervals using a forward-backward search and physiological thresholds. The signal quality index (SQI) is calculated^46^, and the system displays warnings if more than 20% of the RR times series contains RR artifacts (outside of physiological limits or with an SQI below 0.9). A plot displays the filtered ECG signal, identified R-peaks, NN intervals, and interpolated artifacts (see **Figure 4**). NOTE: The RR artifacts are interpolated by default using the shape-preserving piecewise cubic method to obtain the normal-to-normal (NN) intervals, but you can also remove them (not recommended) or use another interpolation method (linear, cubic, nearest neighbor, previous/next neighbor, spline, cubic convolution, or modified Akima cubic). When several ECG channels are present, the RR intervals are estimated for each, and the channel with the least number of RR artifacts is selected for the following steps.
  2.6. Scroll through zoomed-in 30 s windows of the R-peaks for closer inspection by pressing the right/left arrows. If the data contains several ECG/PPG channels, use the channel with the best signal quality index for the RR intervals. BrainBeats does not support both ECG and PPG signals simultaneously at the moment. We chose a sample dataset that has both for tutorial purposes.
  2.7. Once done with the ECG signal, BrainBeats bandpass filters the EEG data at 1-40 Hz using a nonlinear causal minimum-phase FIR filter by default to reduce smearing activity between pre-and post-heartbeat periods, preserve causality and avoid undesired group delays^44^. This is particularly important for users examining the pre-heartbeat period. If the lowpass filter is set to a value above the power line frequency (e.g., 80 Hz lowpass with power line frequency at 50 Hz), use a sharp notch filter to remove the line noise artifact. The EEG data are then re-referenced to infinity using the REST algorithm (best suited for HEP analysis^49^) unless less than 30 channels are detected (in which case they cannot be reliably re-referenced, and a warning is generated to let users know).
  2.8. BrainBeats then detects, removes, and interpolates abnormal EEG channels (**Figure 5**). The default parameters are flat lines greater than 5 s (clean_flatlines algorithm), a maximum high-frequency noise standard deviation of 10, a window length of 5 s (to better capture slow-frequency artifacts^52^), a minimum correlation between neighboring channels of .65, and a maximum tolerated portion of 33% (clean_channels algorithm).
  NOTE: The number of RANSAC samples is set to 500 by default to increase convergence and replicability of bad channel rejection (although it increases the computation time).
  2.9. Next, R-peaks are inserted as event markers into the EEG data to mark each heartbeat, and the data are segmented around these markers without baseline removal (per guidelines^43,66^; **Figure 6**). Since NN intervals have different lengths and EEG must be segmented at a constant length, estimate the minimum epoch size cutoff following R-peak events using the 5th percentile of the interbeat-interval (IBI) data (i.e., the value below which 5% of the shortest IBIs fall, displayed as a dashed red line on a histogram; see **Figure 7**). This 5^th^ percentile value is a good compromise to preserve as many epochs as possible while making sure they are not too short since the period of interest for HEP/HEO analysis is 200-600 ms post heartbeat^49,67^.
  2.10. EEG data are segmented from −300 ms before the R-peaks to the 5^th^ percentile value post-R-peak, with the R-peak at time 0. Epochs shorter than 550 ms or containing more than one R-peak (which would bias the ERP/ERSP) are rejected, per guidelines^49,67^. Epochs containing large EEG artifacts are detected using root-mean-square (RMS) and signal-to-noise ratio (SNR) metrics and MATLAB’s isoutlier function (**Figure 8**). Artifactual epochs are removed.
  2.11. Perform blind source separation using the preconditioned independent component analysis (ICA) algorithm for fast computation^53^ and accounting for data rank for best performance^55^. If desired, choose the ICA Infomax algorithm with special parameters for generating replicable IC maps by choosing the option **Long but replicable** for the field **ICA method** (although this involves longer computation times). Use the ICLabel plugin^56^ to automatically classify ICs to extract non-brain artifacts (ocular components are removed with 90% confidence, whereas muscular, line noise and channel noise are removed with 95% confidence; **Figure 9**).
  2.12. Since **Visualize outputs** is selected in the first GUI window, a scroll plot displays the final signal, and three plots are generated to display: the HEP for each EEG channel (**Figure 10**; with the possibility to click on each channel to look closer), the grand average HEP (**Figure 11 top**) and HEP over time (**Figure 11 bottom**), and the grand average heartbeat-evoked oscillations (HEO; **Figure 12**). HEOs are examined in terms of event-related spectral perturbation (ERSP, i.e., changes in EEG power across heartbeats; **Figure 12 top**) and inter-trial phase coherence (ITC; i.e., consistency of the phase angle across heartbeats; **Figure 12 bottom**). NOTE: ERSP is computed using a default 3-cycle wavelet (with a Hanning-tapered window applied, pad ratio of 2) and with the number of cycles in the wavelets used for higher frequencies expanding slowly up to 20% of the number of cycles in the equivalent FFT window at its highest frequency (1 minus 0.8). This controls the shapes of the individual windows measured by the function and their shapes in the resulting time/frequency panes. An arbitrary baseline is removed for illustration purposes, and ERSP is computed for frequencies 7-25 Hz to capture the typical HEO effect described in the literature, namely 300-450 ms post heartbeat in the alpha band (8-13 Hz) over frontocentral electrodes^17,67^. Lower frequencies cannot be estimated due to the short epoch size defined by the interbeat intervals. Nonparametric (permutation) statistics are applied to visualize the HEO for a p-value of 0.05, corrected for false discovery rate (FDR; i.e., type 1 error or family-wise error). These plots are generated for tutorial purposes or single-trials analysis.
  2.13. Preprocessing plots are generated to visualize the different steps. To turn it off, uncheck the box **Visualize preprocessings (Figure 2)**. The final EEG data (cleaned and segmented around the R-peaks) do not include the ECG data since it would bias the ERP/ERSP analysis. To preserve the heart channel in the final output, check the box **Keep heart channel**.
  2.14. You can pause before processing the next file (next condition or participant). Finally, BrainBeats supports EEGLAB’s history function. At the end of all operations, type **eegh** in MATLAB’s command window to print the command line to repeat all the above steps via a single command line, with the parameters that were selected manually in the GUI, allowing easy automation. You can find preprocessing outputs (e.g., signal quality index of the cardiovascular time series, NN intervals, removed EEG channels, segments, and components, etc.) in the EEGLAB structure: EEG.brainbeats.preprocessings. All parameters are also exported in EEG.brainbeats.parameters. We encourage users to report these outputs in scientific publications to increase the replicability of findings.
  2.15. For advanced users, perform all the above steps with default parameters with the following command lines (see the tutorial script in the BrainBeats repository for more options): eeglab; close; % Launch EEGLAB without the GUI main_path = fileparts(which(’eegplugin_BrainBeats.m’)); cd(main_path); EEG = pop_loadset(‘filename’,‘ dataset.set’, ‘filepath’,fullfile(main_path, ‘sample_data’)); % Load the sample dataset EEG = brainbeats_process(EEG,‘ analysis’, ‘hep’, ‘heart_signal’, ‘ECG’, … ‘heart_channels’,{‘ECG’}, ‘clean_eeg’,true); % Launch BrainBeats 1^st^ level to process the file for HEP analysis with default parameters
  2.16. The steps above performed HEP/HEO from the ECG signal. Use the following steps for the PPG signal.
  2.17. In the following steps, perform the same operations but using PPG signal. Load the same dataset again (see step 1.5) since the previous operations overwrote it, and open BrainBeats’ first GUI again to select the main parameters. Click **Tools > BrainBeats > 1^st^ level (subject level).** Select **Heartbeat-evoked potentials (HEP)** as the analysis to run, **PPG** as the heart data type, and click on the button to display the list of channels to select the PPG channel. Keep the options **Visualize outputs (recommended)** and **Save outputs** selected and click **Ok**. See **Figure 13**.
  2.18. The second GUI window pops up in a manner similar to step 2.2. Except for the parameter options for processing the cardiovascular signals that are now adapted to the PPG signal (**Figure 14**). Click **Ok** to run with the default parameters.
  2.19. A warning message appears, asking for confirmation to remove the extra ECG channel that has been detected. Again, this is expected. Click **Yes**. By default, the toolbox will preprocess the PPG signal, detect the pulse waves to obtain the RR intervals, identify RR artifacts, if any, and interpolate them (**Figure 15**). Steps 2.7. to 2.12. are performed and the same plots and outputs are generated but based on the R waves detected from the PPG signal (see **Figures 16, 17, and 18**). NOTE: R waves are detected using the slope of the signal within a specified window. Potential pulses are then flagged when the slope exceeds a dynamic threshold, which is adjusted based on the detection history and signal characteristics. The algorithm then searches within an “eye-closing” period to pinpoint the maximum slope, and subsequently, the onset of the pulse wave is determined through thresholding. The R-wave peaks are identified as the valleys near the onset, and their locations are recorded. The algorithm iterates through the entire signal, continuously adjusting detection thresholds and identifying R-wave peaks, which are then used to calculate the RR intervals.
  2.20. For advanced users, perform all the above steps with default parameters with the following command lines (see the tutorial script in the BrainBeats repository for more options): eeglab; close; % Launch EEGLAB without the GUI main_path = fileparts(which(‘eegplugin_BrainBeats.m’)); cd(main_path); EEG = pop_loadset(‘filename’,‘ dataset.set’, ‘filepath’,fullfile(main_path, ‘sample_data’)); % Load the sample dataset EEG = brainbeats_process(EEG,‘ analysis’, ‘hep’, ‘heart_signal’, ‘PPG’, … ‘heart_channels’,{‘PPG’},‘clean_eeg’,true); % Launch BrainBeats 1^st^ level to process the file for HEP analysis with default parameters
**3. Extract EEG and HRV features**
  3.1 Load the same dataset again (see step 1.5; Click **File** > **Load existing dataset** > **Select dataset.set**) since it was overwritten by the previous operations and open the main GUI again to select the main parameters (step 2.1; Click **Tools > BrainBeats > 1^st^ level**; see **Figure 1**). Select **Extract EEG & HRV features** for the analysis type, **ECG** for the heart signal type, and select **ECG** in the list of electrode labels. Click **Ok**.
  3.2 The second GUI window pops up like in step 2.2 but with different parameters for EEG preprocessing and extracting HRV and EEG features (**Figure 19**). In the HRV section, click on the button **freq. options** to select the method to compute HRV power (default set to normalized Lomb-Scargle periodogram), the window overlap (default set to 25%), and for performing a second-level normalization (unset by default; see note below for more detail). Similarly, in the EEG features section, click on the button **freq. options** to fine-tune some parameters, such as the overall frequency range on which to compute the power spectral density (PSD; default = 1-40 Hz), the units (decibels, µV^2^/Hz, or normalized by total power), the window type (default = hamming), the window overlap (default = 50%), the window length (default = 2 s), and the types of frequency bounds for each band. Click **Ok** to launch with default parameters. NOTE: HRV power is computed by default using the normalized Lomb-Scargle periodogram, which does not require resampling (therefore better preserving original information) and best deals with non-uniformly sampled data, missing data, and noise (typical with NN intervals)^68^. The normalized version scales the power by the variance of the signal, providing results that are less sensitive to varying noise levels, more focused on the relative strength of periodic components, and more comparable across different recordings or subjects. Other methods available include the non-normalized Lomb-Scargle periodogram, the Welch method, and the Fast Fourier Transform (FFT). Resampling is automatically performed for the Welch and FFT methods to create the necessary regularly sampled time series. A second-level normalization can be applied by dividing the power of each frequency band by the total power, providing a more intuitive measure of the relative contribution of each frequency component to the overall power. It is disabled by default since it is only meaningful when all four bands are available, requiring at least 24 h of signal. These algorithms are adapted from the Physionet Cardiovascular signal processing toolbox^46^.
  3.3 The same warning message appears, asking for confirmation to remove the extra PPG channel that has been detected. Again, this is expected. Click **Yes**. BrainBeats will start preprocessing the ECG data and extracting the NN intervals identically, as in step 2.4. Then, it extracts heart-rate variability (HRV) features from the NN intervals in the time (SDNN, RMSSD, pNN50), frequency (ULF, VLF, LF, HF, LF:HF ratio, total power), and nonlinear (Poincare, phase-rectified signal averaging, fuzzy entropy, and fractal dimension) domains. NOTE: BrainBeats automatically checks the file length to ensure that minimum requirements are met (e.g., ULF-HRV power requires 24 hours of data), sends warning messages if not, and does not export these features to prevent unreliable estimations. BrainBeats follows guidelines and recommendations for estimating HRV metrics^27,45^.
  3.4 Next, BrainBeats preprocesses the EEG data as in step 2.7**. (see Figure 5**). Then, large artifacts are detected automatically in the continuous data using the artifact subspace reconstruction (ASR) algorithm^50,52^ (default SD criterion set to 30 and using 80% of available RAM to increase speed). These large artifacts are removed from the EEG data (see **Figure 20**). To adjust these parameters in the GUI, select the fields **Threshold to reject bad segments with ASR** and **Available RAM to use for ASR**. NOTE: The EEG and cardiovascular time series do not need to be time synchronized for the features mode since the features are estimated on each signal separately. Thus, EEG artifacts can be removed directly from the EEG data (in red, **Figure 20**), unlike for the HEP mode, where epochs containing artifacts were rejected for both time series because time synchronization with ms accuracy is essential for that method.
  3.5 Then, perform ICA using the same algorithms and parameters as for HEP (see step 2.11.), except that this time the heart components are removed if detected with 99% confidence (they were preserved for HEP/HEO since we do not want to remove relevant cardiac-related signals).
  3.6 Because you checked the box **Frequency domain** in step 3.2., BrainBeats extracts the following frequency domain features: the average power spectral density (PSD) for the delta (1-3 Hz), theta (3-7 Hz), alpha (8-13 Hz), beta (13-30 Hz), and gamma (30+ Hz) frequency bands, the individual alpha frequency (IAF), and alpha asymmetry on all available (symmetric) electrode pairs. NOTE: PSD converted to decibels (dB) facilitates the comparison of results across recordings and subjects. The frequency bounds can be set to the conventional bounds (e.g., predefined 8-13 Hz for the alpha band) or to the individualized bounds, which are detected from the distribution of the power spectral density to account for inter-individual differences^69^ (e.g., 7.3-12.6 Hz for the alpha band). The algorithm was designed for the alpha band and does not perform as well for other bands, especially when peaks are not present in the power spectral distribution. The individual alpha frequency (IAF) is estimated using the alpha center of gravity to better deal with split peaks or ambiguous peaks^69^. Alpha asymmetry is computed following guidelines (2-s hamming window with 50% overlap, the logarithm of alpha power from the left channel minus logarithm of alpha power from the right channel)^47^. Hence, positive values indicate greater left-than-right alpha power and vice versa. Alpha asymmetry can be normalized by dividing alpha power from each electrode by the alpha power summed across all electrodes^47^. The symmetric pairs are obtained using theta distances, requiring the EEG data to contain electrode coordinates.
  3.7 Because you checked the box **Nonlinear domain** in step 3.2., BrainBeats extracts the fuzzy entropy and the fractal dimension for each EEG channel. NOTE: Nonlinear-domain features are thought to capture nonlinear, complex dynamics of the brain that are missed by spectral measures and show particular promise for investigating interactions between various body systems ^70–72^. Fuzzy entropy is more reliable and robust than its alternatives (sample and approximate entropy) but requires longer computation times (especially with long EEG time series with high sampling rates). To address this issue, when EEG signals are longer than 2 minutes long with a sampling rate greater than 100 Hz, they are automatically downsampled (or decimated when the factor is not an integer) to 90 Hz (i.e., corresponding to a Nyquist frequency of 45 Hz, to match the default lowpass filter and avoid line noise artifacts as much as possible). Furthermore, parallel computing is activated by default when estimating EEG features, which reduces computing time, especially when many EEG channels are available.
  3.8 Because **Visualize outputs** was selected in the first GUI (see step 3.1. and **Figure 1**), BrainBeats generates a plot displaying the power spectral density (PSD) for HRV and EEG data (**Figure 21**) along with scalp topographies displaying some EEG features (**Figure 22**). NOTE: You can also find some preprocessing outputs in EEG.brainbeats.preprocessing and all the parameters used in EEG.brainbeats.parameters. We encourage users to report these outputs in scientific publications to increase the replicability of findings.
  3.9 Keep the **Save outputs** box checked in the first GUI window to save All features are exported into the EEGLAB .set file in EEG.brainbeats.features and saved in a .mat file in the same folder where the dataset was loaded.
  3.10 Finally, BrainBeats supports EEGLAB’s history function. At the end of all operations, type **eegh** in MATLAB’s command window to print the command line that will allow you to repeat all the above steps via a single command line, with the parameters that were selected manually in the GUI, allowing easy automation and replication of the operations.
  3.11 For advanced users, perform all the steps above with the following command: eeglab; close; % Launch EEGLAB without the GUI main_path = fileparts(which(‘eegplugin_BrainBeats.m’)); cd(main_path); EEG = pop_loadset(‘filename’,‘ dataset.set’, ‘filepath’,fullfile(main_path, ‘sample_data’)); % Load the sample dataset EEG = brainbeats_process(EEG,‘ analysis’, ‘features’, ‘heart_signal’, ‘ECG’, … ‘heart_channels’,{‘ECG’},‘clean_eeg’,true);
  3.12 The previous steps extracted HRV features from ECG signal. Use the following steps to extract HRV features from PPG signal (EEG features are the same).
  3.13 Load the same dataset again (step 1.5.) since it was overwritten by the operations and open the main GUI again (step 2.5.). Select **Extract EEG & HRV features** for the analysis and select **PPG** for the heart signal type and **PPG** for the channel name. Click **Ok**.
  3.14 The 2^nd^ GUI window now shows the parameters for preprocessing PPG and for extracting the HRV and EEG features. Click **Ok** to run with the default parameters. Parameters are described in step 2.17.
  3.15 A warning message will appear, asking for confirmation to remove the detected ECG channel. This is to be expected since the toolbox is not designed to analyze both ECG and PPG simultaneously (or other auxiliary channels) and keeping it in the dataset will lead to serious errors (e.g., artifact removal, poor ICA decomposition, etc.). Click **Yes**.
  3.16 BrainBeats preprocesses the PPG signal and estimates the NN intervals as in step 2.5. It then extracts the HRV features from the NN intervals, the same as in step 3.2. except that the NN intervals have now been obtained from the PPG signal. The EEG signals are preprocessed as in step 3.2. BrainBeats plots the PSD (**Figure 23).** The only difference here is the PSD estimated from the NN intervals obtained from the PPG as opposed to ECG.
  3.17 For advanced users, perform all the steps above with the following command: eeglab; close; % Launch EEGLAB without the GUI main_path = fileparts(which(‘eegplugin_BrainBeats.m’)); cd(main_path); EEG = pop_loadset(‘filename’,‘ dataset.set’, ‘filepath’,fullfile(main_path, ‘sample_data’)); % Load the sample dataset EEG = brainbeats_process(EEG,‘ analysis’, ‘features’, ‘heart_signal’, ‘PPG’, … ‘heart_channels’,{‘PPG’},‘clean_eeg’,true);
**4. Extract heart artifacts from EEG signals.**
  4.1 Load the sample dataset (see step 1.5.).
  4.2 Open the main GUI window by clicking **Tools > BrainBeats > 1^st^ level (subject level)** (**Figure 1**) and select **Extract heart artifacts from EEG signals** for the analysis type, **ECG** for heart signal type and select **ECG** in the list of electrode labels. Click **Ok**. See **Figure 24**.
  4.3 The 2^nd^ GUI window (**Figure 25**) shows the preprocessing parameters. Set the **Power line noise** to **50 Hz (Europe),** edit the confidence level if needed, check the box **Boost mode (beta)**, and click **Ok** to run with the default parameters since the EEG signals from the sample dataset are not preprocessed. NOTE: the confidence level to detect heart components is set to 80% by default, which may be too low or too high for some datasets. Increasing this value will increase the chances of detecting heart components but decrease the reliability of that detection. The Boost mode (beta) is optional and aims to improve classification performance by smearing the heart signal into the EEG signals.
  4.4 A warning message will appear, asking for confirmation to remove the extra PPG channel that has been detected. This is to be expected since the toolbox is not designed to analyze both ECG and PPG simultaneously (or other auxiliary channels) and keeping it in the dataset will lead to serious errors (e.g., artifact removal, poor ICA decomposition, etc.). Click **Yes**.
  4.5 The ECG signal is bandpass-filtered to remove slow frequency drifts below 1 Hz and high-frequency noise above 20 Hz (with a noncausal zero-phase FIR filter). The EEG signals are preprocessed as in step 3.4.
  4.6 Independent component analysis (ICA) is performed using the preconditioned ICA for real data algorithm (PICARD). To change this option, choose the standard Infomax algorithm or modified Infomax algorithm for replication from the GUI in step 4.3. Then, classify the independent components automatically with ICLabel. If a component is classified as a heart component with 80% confidence, it is, by default, detected and extracted automatically from the EEG data.
  4.7 Keep the box Visualize outputs in the first main GUI (step 4.2. and **Figure 24**) to visualize the scalp topography of the removed component(s) (**Figure 26**) and the final EEG time series (in blue; **Figure 27**) after extracting the heart component(s) (in red; **Figure 27**). NOTE: the ECG channel is preserved for visualization to confirm the extraction of ECG-related components, but it is removed after this step since it no longer contains any relevant information.
  4.8 For advanced users, perform these steps using the following command lines: For advanced users, perform these steps using the following command lines: eeglab; close; % Launch EEGLAB without the GUI

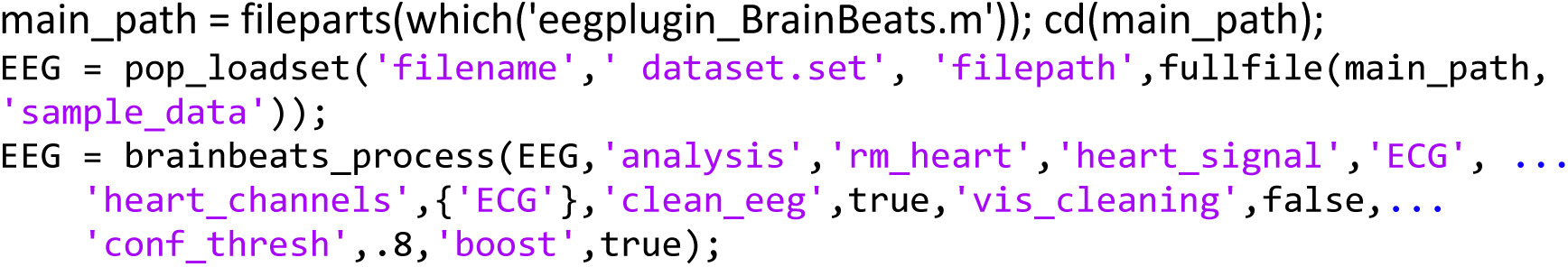

**FIGURE 1.**
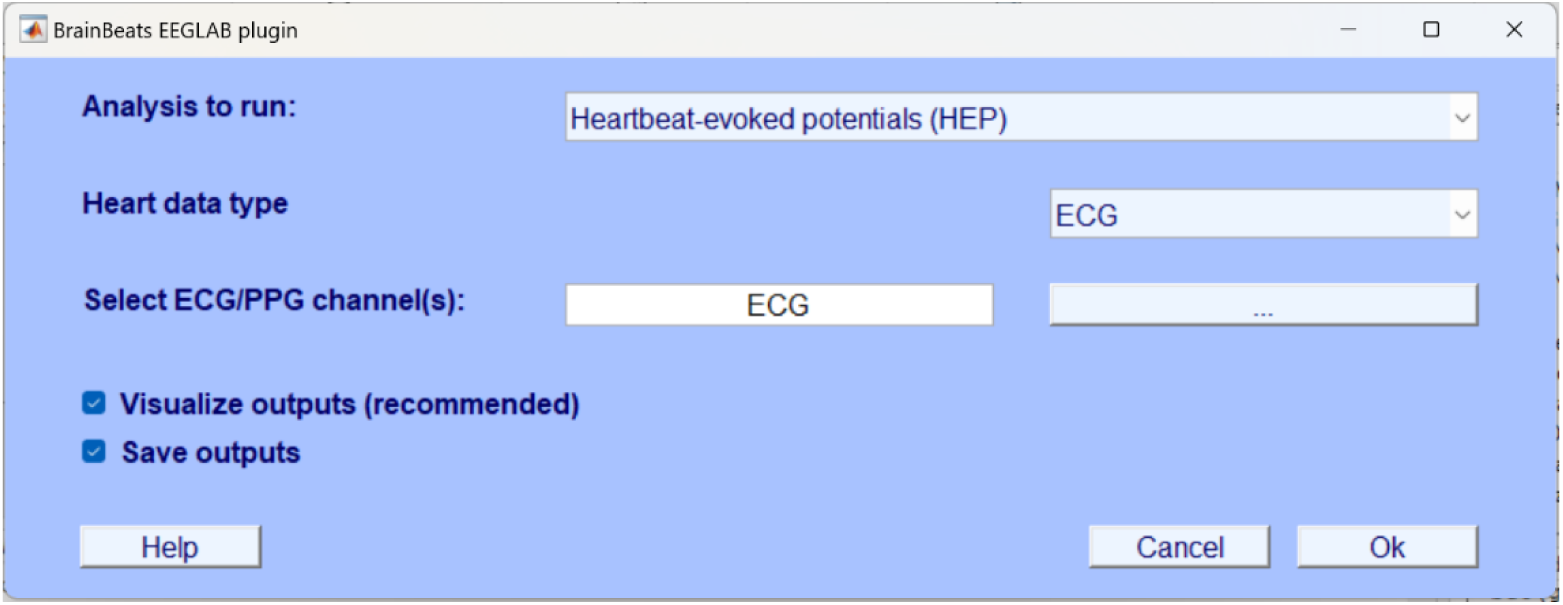
BrainBeats’ main menu (1^st^ level). Select general parameters from the main general user interface (GUI) to process a file for HEP/HEO analysis.

**FIGURE 2.**
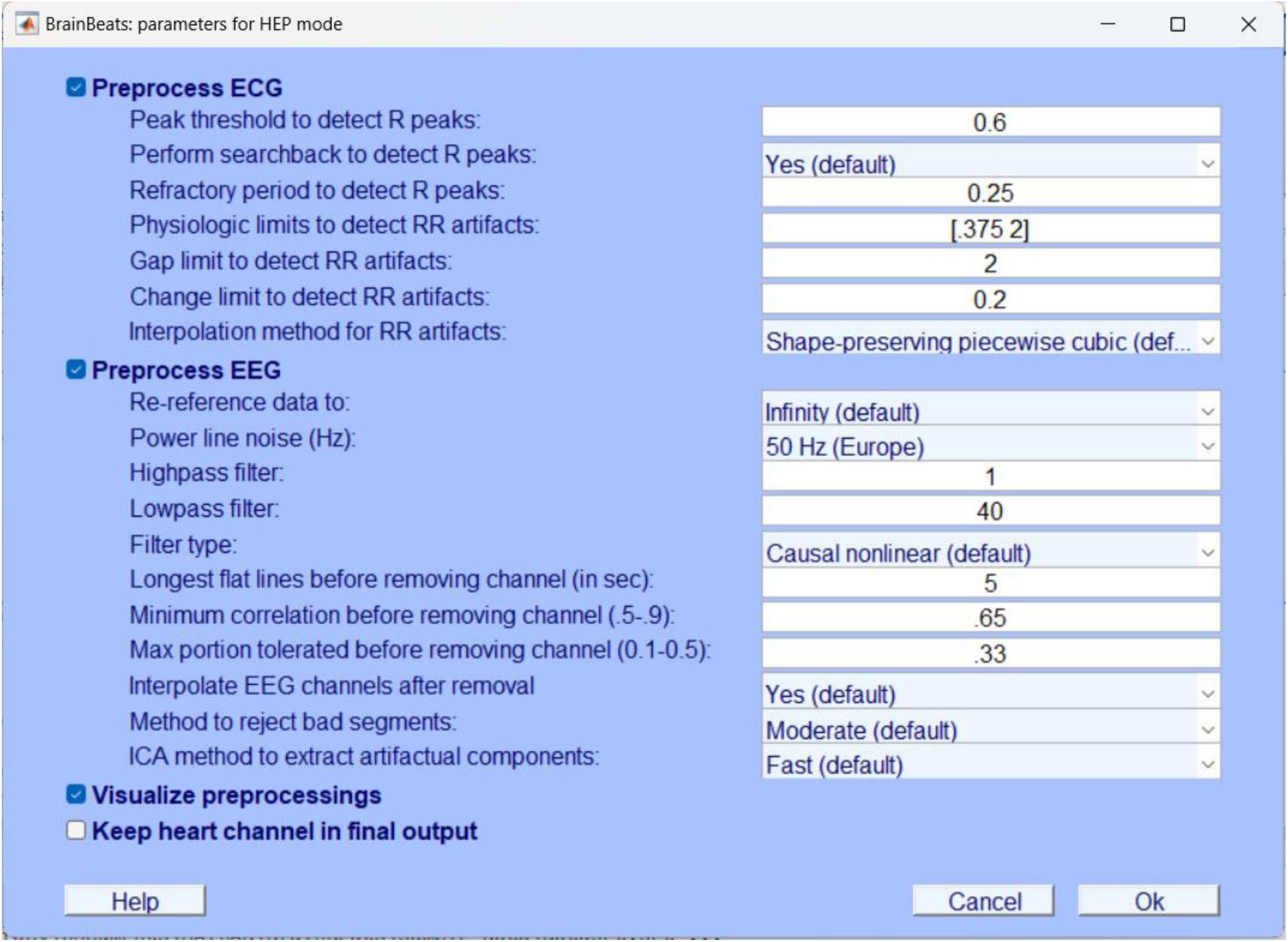
Fine-tuning parameters for preprocessing ECG and EEG signals for HEP/HEO analysis. This graphic user interface (GUI) allows users to fine-tune the main preprocessing parameters to: 1) obtain the RR intervals from ECG signals, detect and interpolate RR artifacts if any to obtain the NN intervals; 2) filter and re-reference EEG data, detect and interpolate abnormal EEG channels if any, extract non-brain artifacts using ICA; 3) choose if wish to visualize preprocessing steps (for fine-tuning), or keep the cardiovascular channel in the final output (not recommended for HEP/HEO).

**FIGURE 3.**
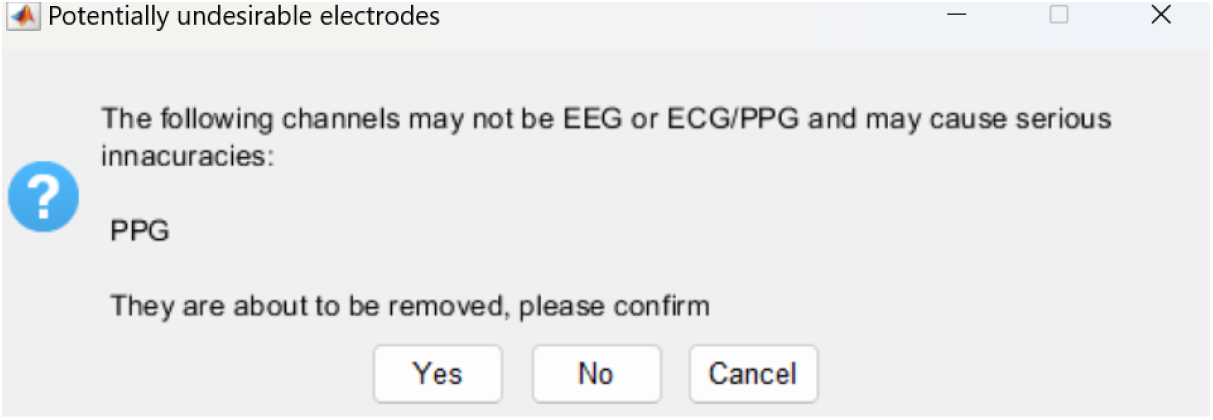
Extra channel warning. A warning message asks for user confirmation to remove the extra PPG channel detected in this dataset. This is to be expected because the toolbox is not currently designed to process both ECG and PPG signals simultaneously.

**FIGURE 4.**
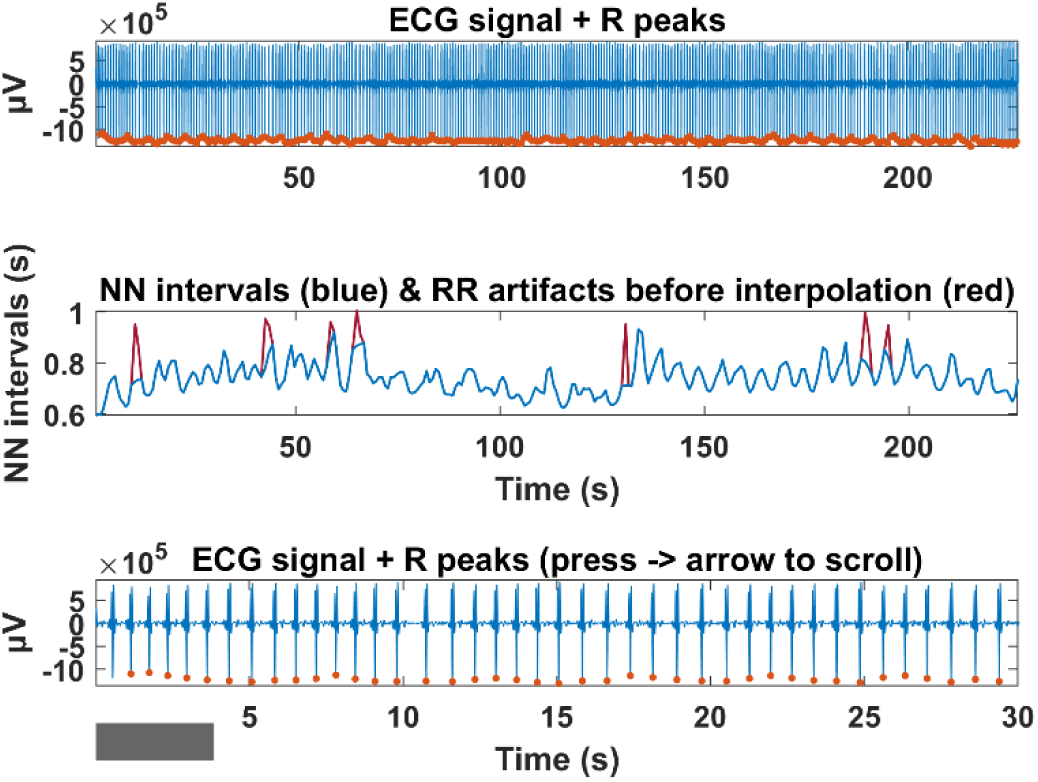
RR intervals, artifacts, and NN intervals obtained from ECG signal. **Top**: preprocessed ECG signal (blue) with the R-peaks detected by BrainBeats (orange dots, i.e. RR intervals). **Middle**: Normal-to-Normal (NN) intervals (in blue) after interpolation of the RR artifacts (in red). **Bottom**: Same as top plot but zoomed in (30-s window) to inspect the R-peaks more closely with a scrolling feature by pressing the left/right arrows.

**FIGURE 5.**
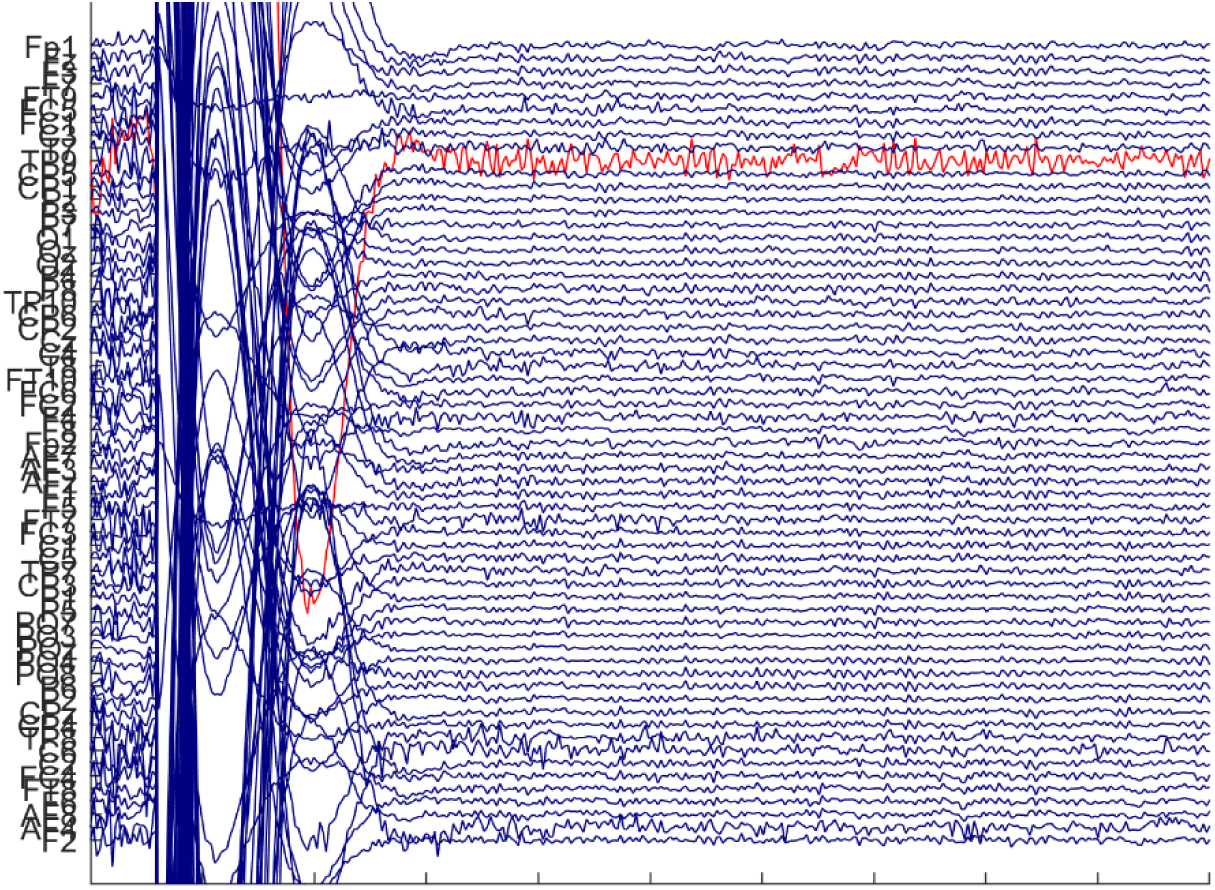
Rejection of bad EEG channels. Visualization of the abnormal EEG channel (TP9) automatically detected and removed from the dataset. Note: the large artifact is dealt with in a further step. EEG data were bandpass filtered (1-40 Hz) and re-referenced to infinity.

**FIGURE 6.**
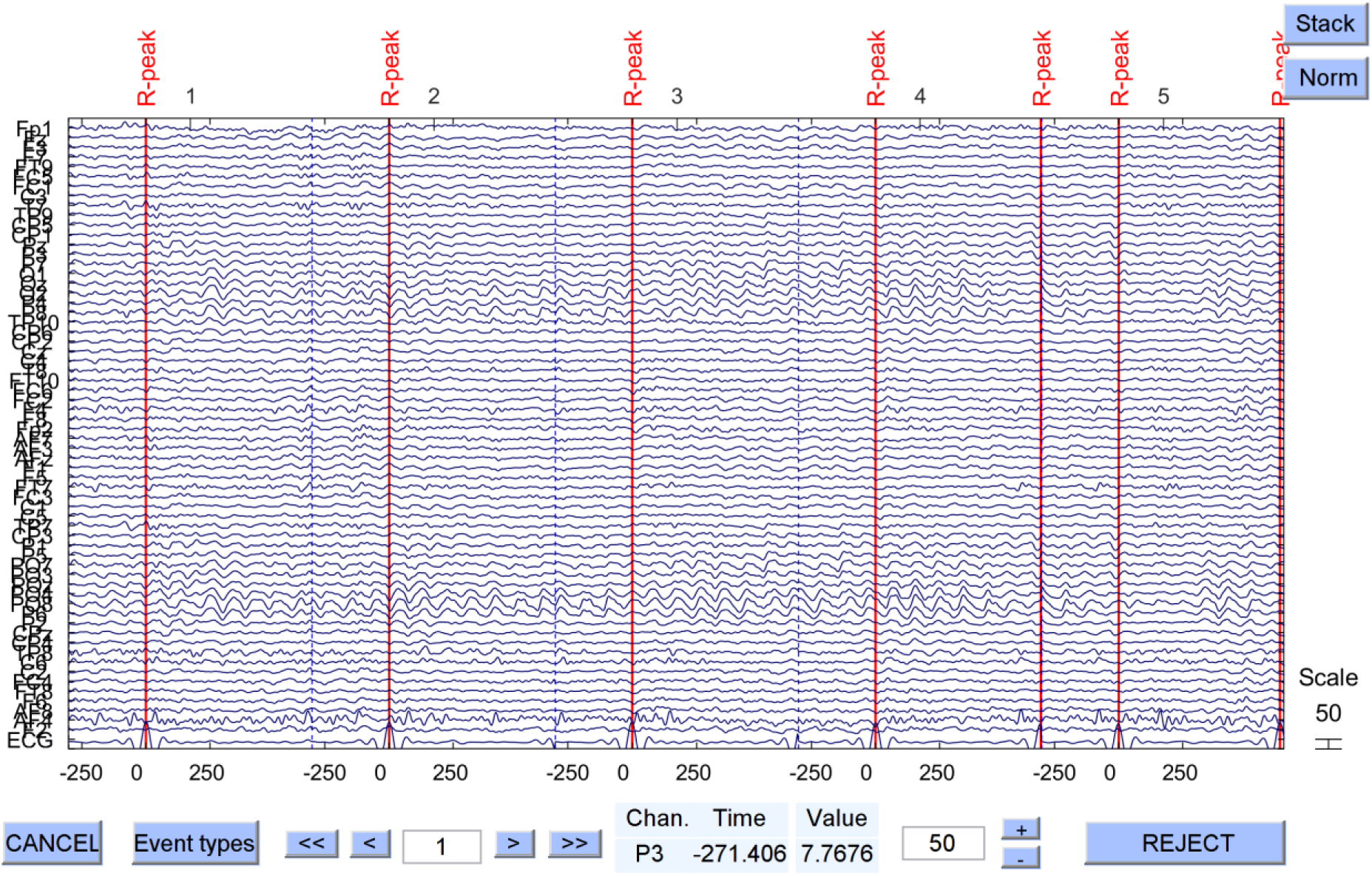
Visualization of the 64-channel EEG data after preprocessing and marking the R-peaks in the signal. The ECG signal is included in the plot at the bottom for visual confirmation.

**Figure 7.**
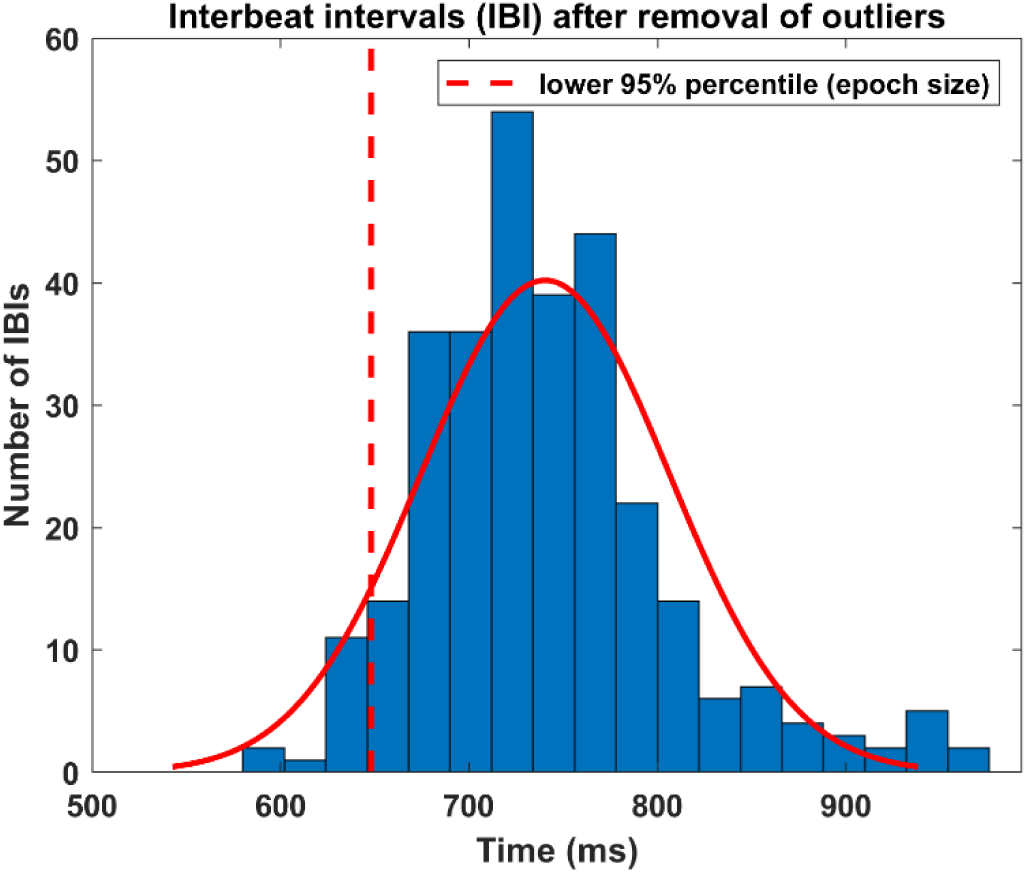
Histogram of the interbeat intervals (IBI). The red line shows the fitted normal distribution and the red dashed line shows the 5% percentile, used as the upper cutoff value at which the EEG data are segmented (i.e., 650 ms after the R-peak here).

**Figure 8.**
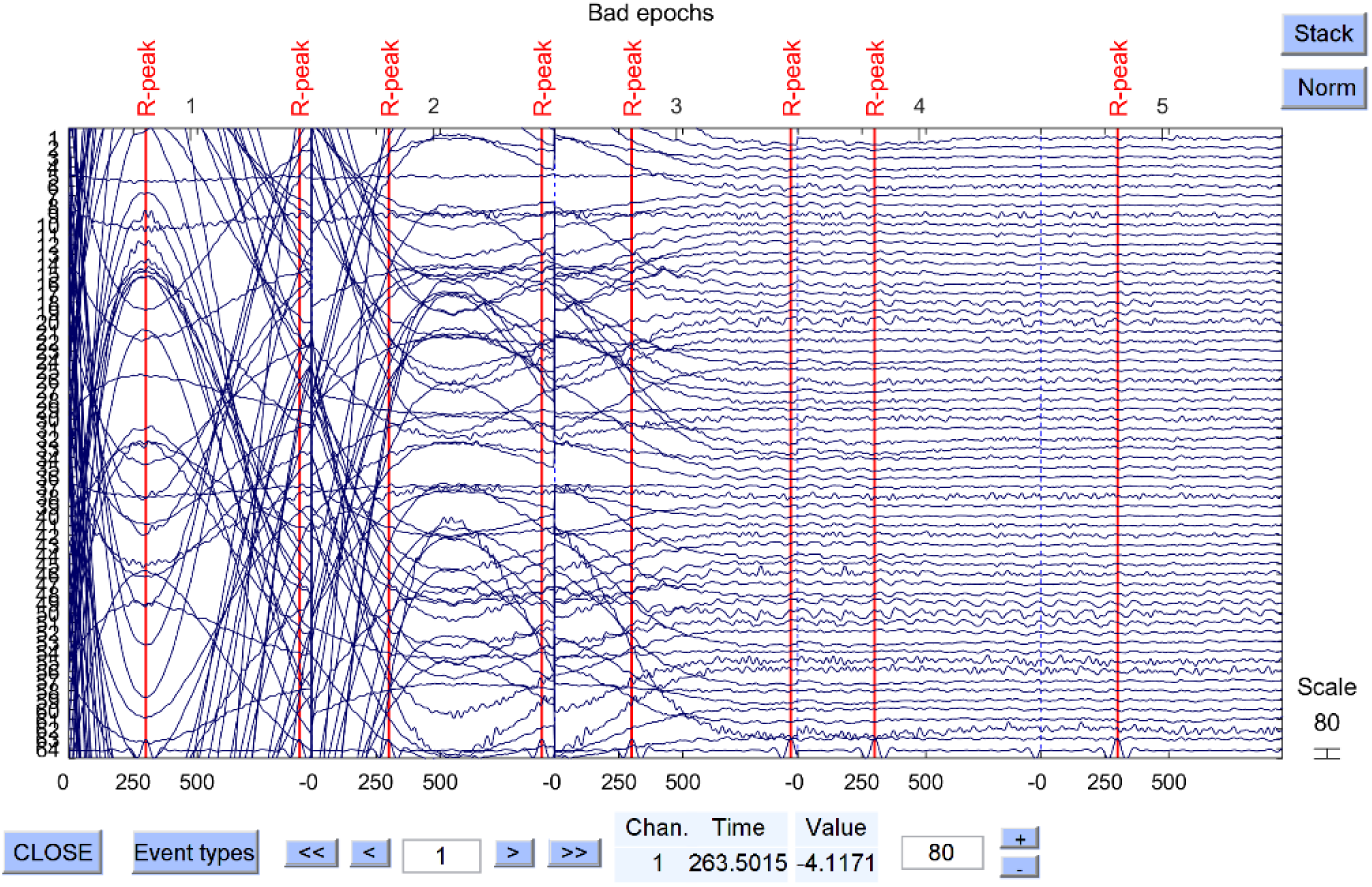
Removal of artifactual epochs. Visualization of the EEG outlier epochs (i.e., containing artifacts) that were detected and removed prior to performing independent component analysis (ICA).

**Figure 9.**
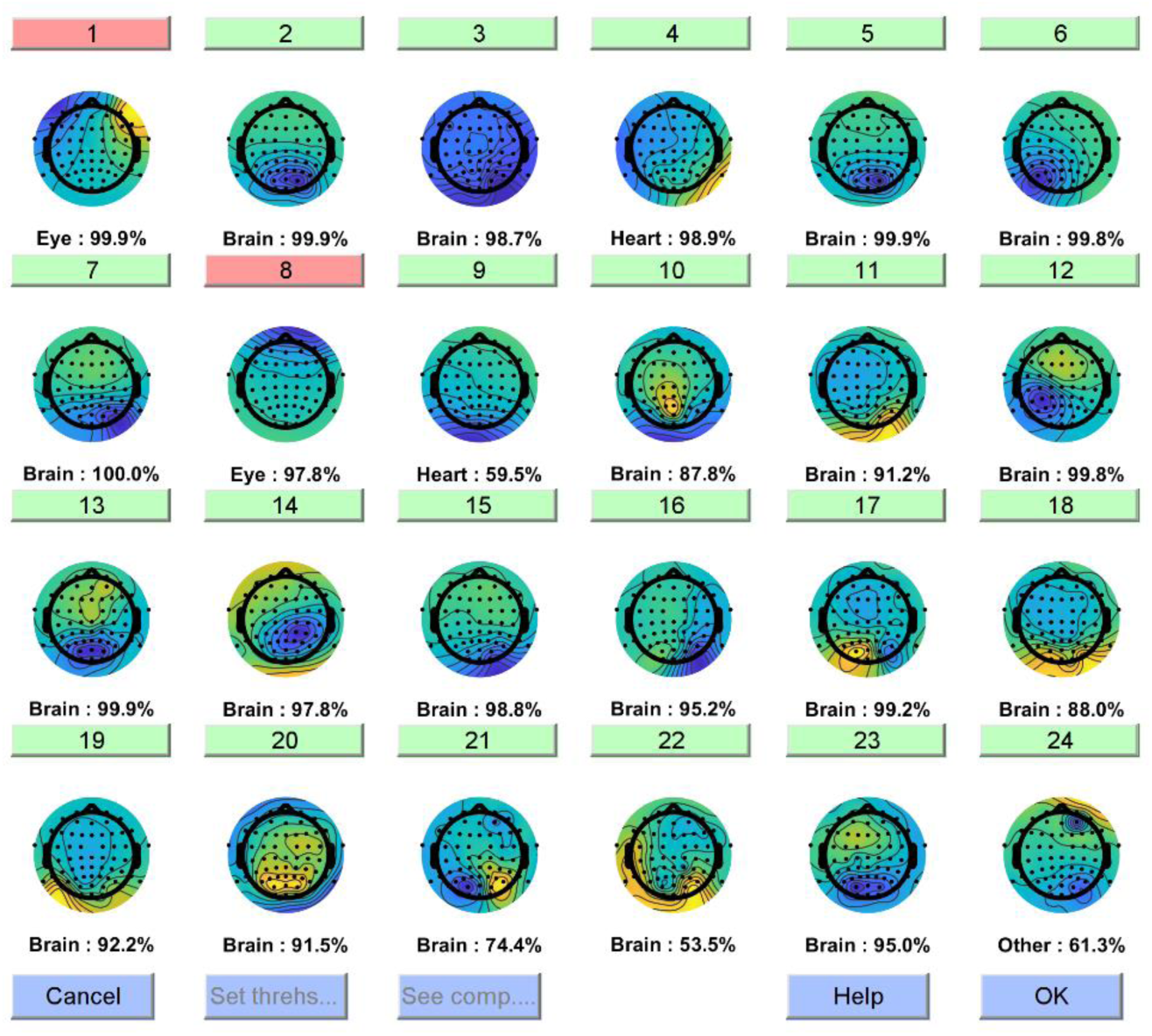
Classification of the independent components to remove non-brain artifacts. After performing blind source separation to obtain the independent components of our EEG data, the ICLabel plugin is used to classify them and automatically flag the non-brain components for extraction.

**Figure 10.**
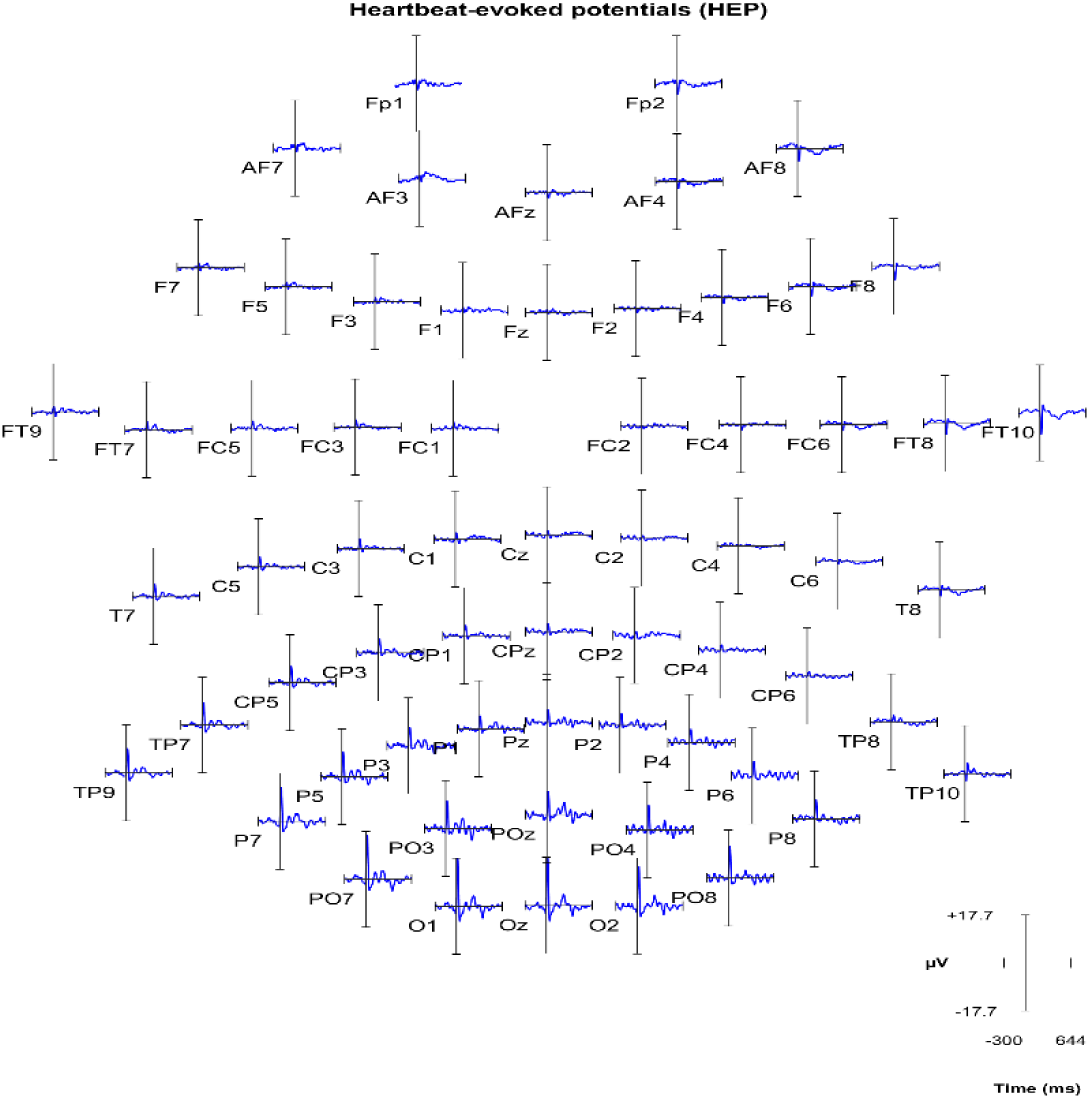
Visualization of the heartbeat-evoked potentials (HEP) for each EEG channel obtained from the ECG signal. Users can click on each channel to inspect it more closely.

**Figure 11.**
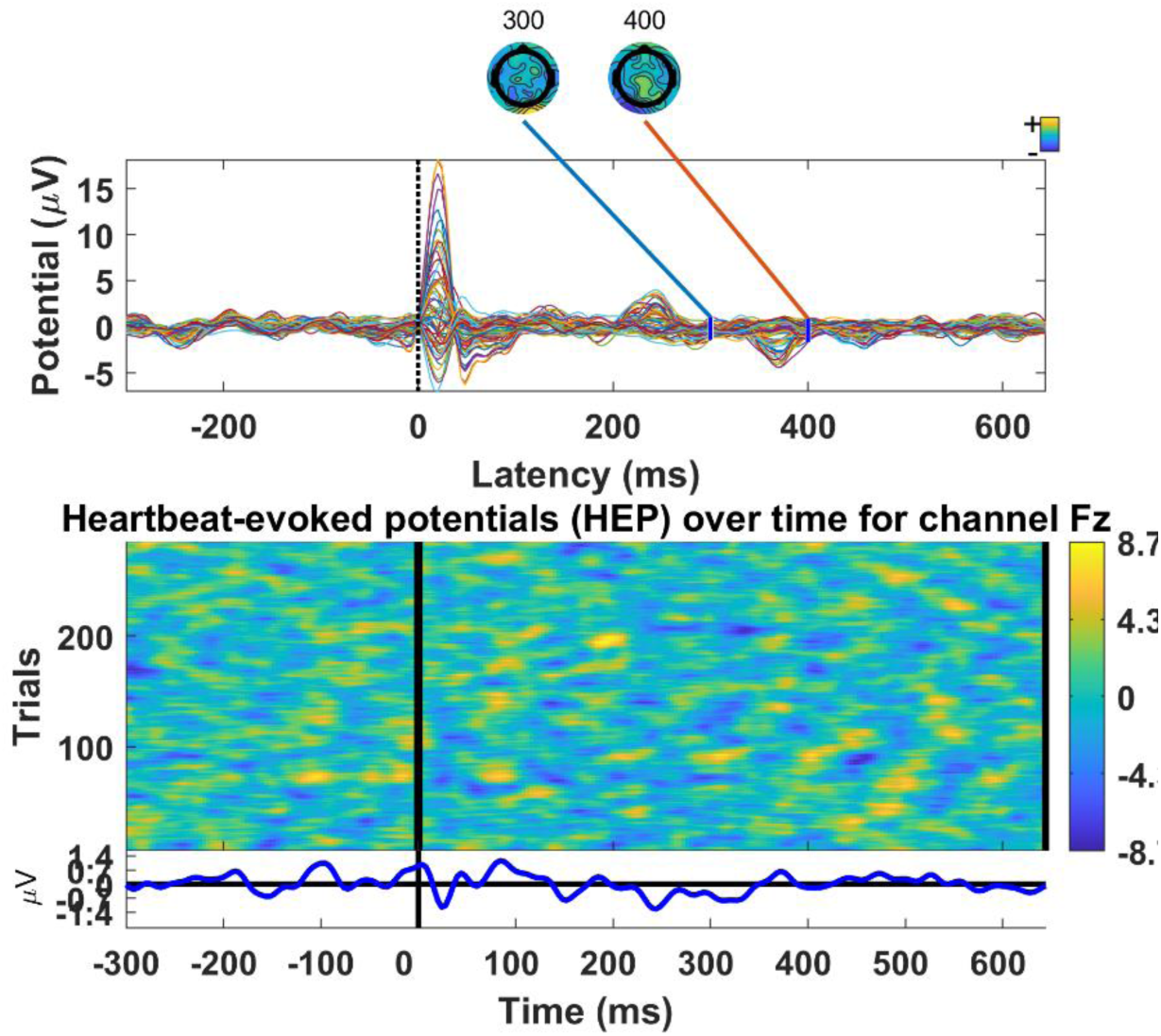
Grand average heartbeat-evoked potentials (HEP) obtained with ECG. **Top**: Average across epochs for each EEG channel (superimposed), with scalp topographies showing amplitude distribution in the period of interest (200-500 ms post R-peak). **Bottom**: HEP evolution over time (each “trial” corresponds to a heartbeat).

**Figure 12.**
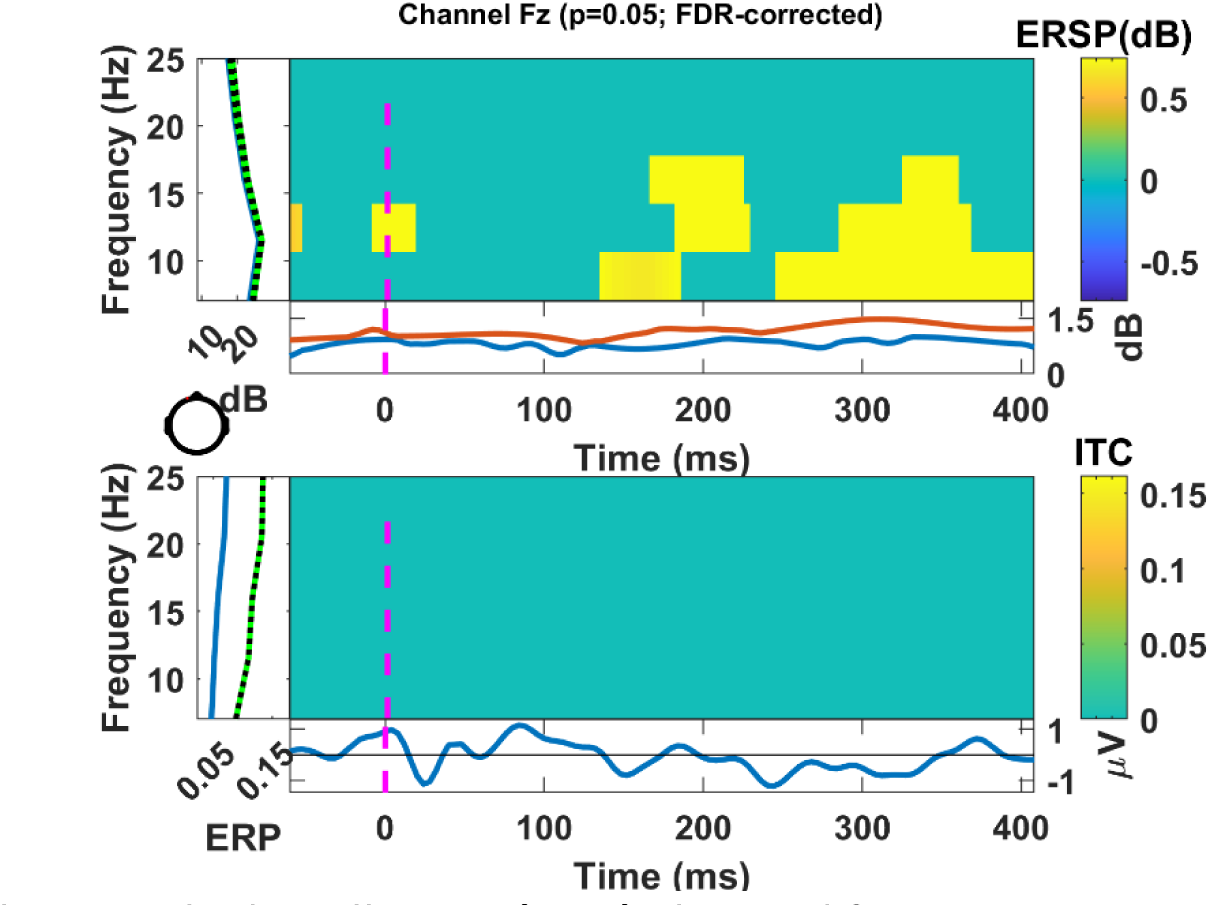
Heartbeat-evoked oscillations (HEO) obtained from ECG. **Top:** HEO at channel Fz (frontocentral region) after permutation statistics (1000-iterations) and corrected for false discovery rate (FDR) at the 95% confidence level (p<0.05). **Bottom**: Inter-trial coherence (ITC) after FDR correction.

**Figure 13.**
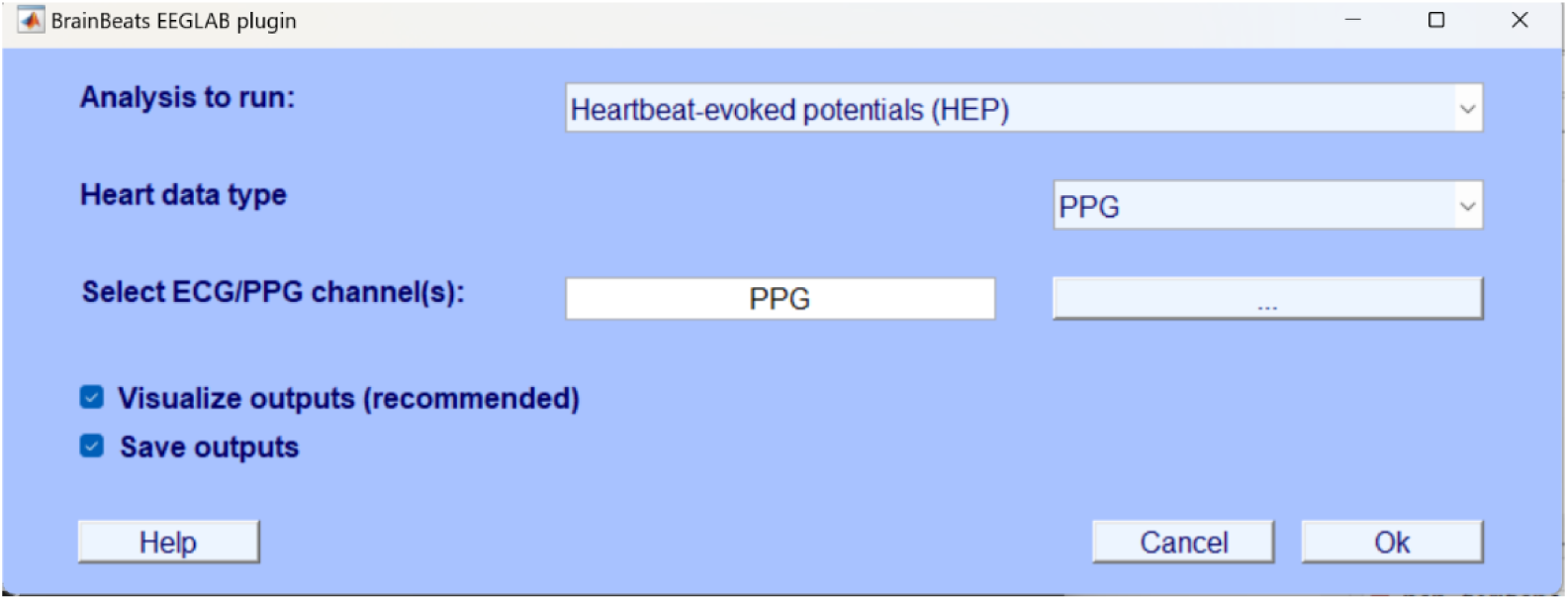
Main GUI window with the general parameters to perform HEP/HEO analysis with PPG signals.

**Figure 14.**
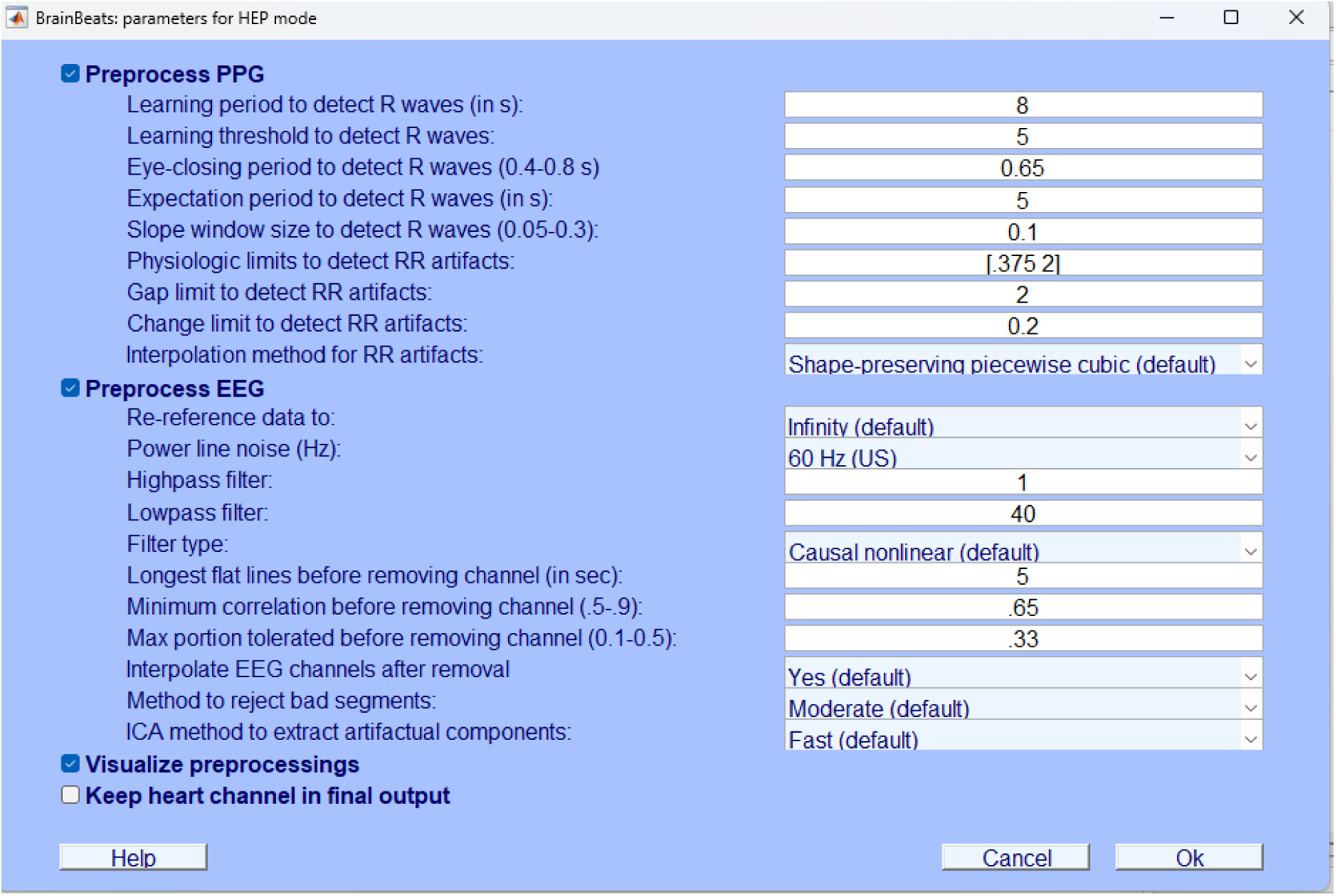
Fine-tuning parameters for preprocessing PPG and EEG signals for HEP/HEO analysis. The graphical user interface (GUI) is the same as Figure 2 for EEG but has different parameters for estimating the RR intervals from the PPG signals.

**FIGURE 15.**
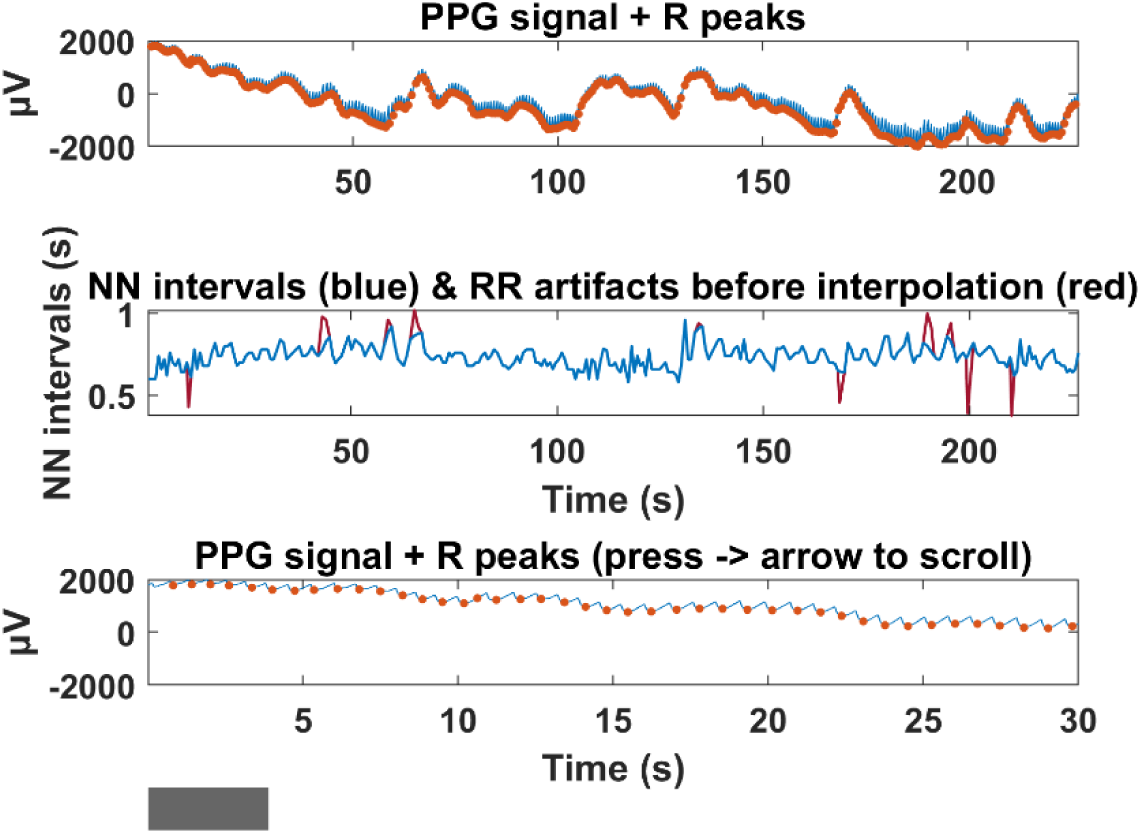
RR intervals, artifacts, and NN intervals obtained from PPG signal. **Top**: preprocessed PPG signal (blue) with the pulse waves detected by BrainBeats (orange dots, i.e. RR intervals). **Middle**: Normal-to-Normal (NN) intervals (in blue) after interpolation of the RR artifacts (in red). **Bottom**: Same as top plot but zoomed in (30-s window) to inspect the pulse waves more closely with a scrolling feature by pressing the left/right arrows.

**Figure 16.**
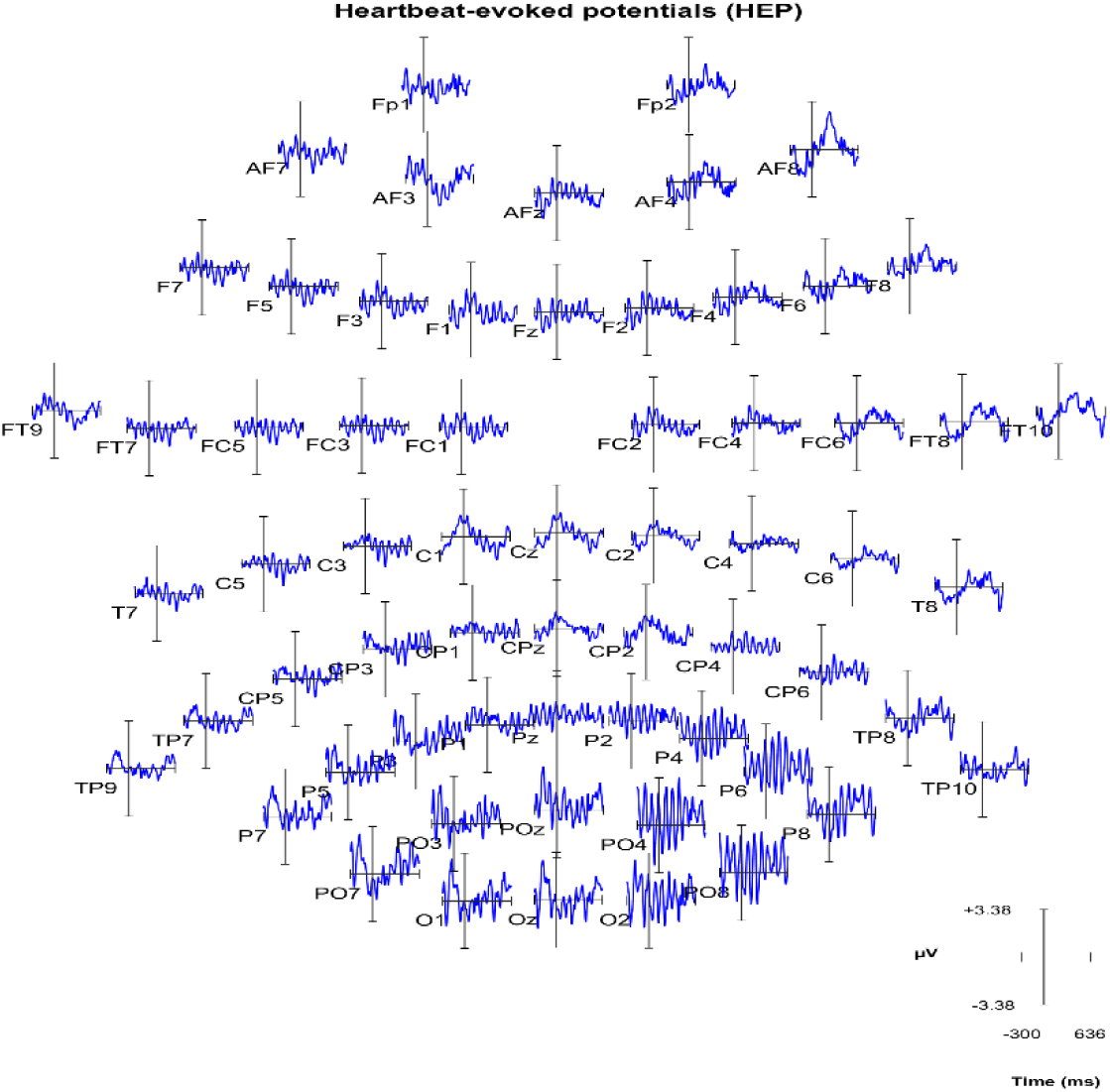
Visualization of the heartbeat-evoked potentials (HEP) obtained from the PPG signal for each EEG channel. Users can click on each channel to inspect it more closely.

**Figure 17.**
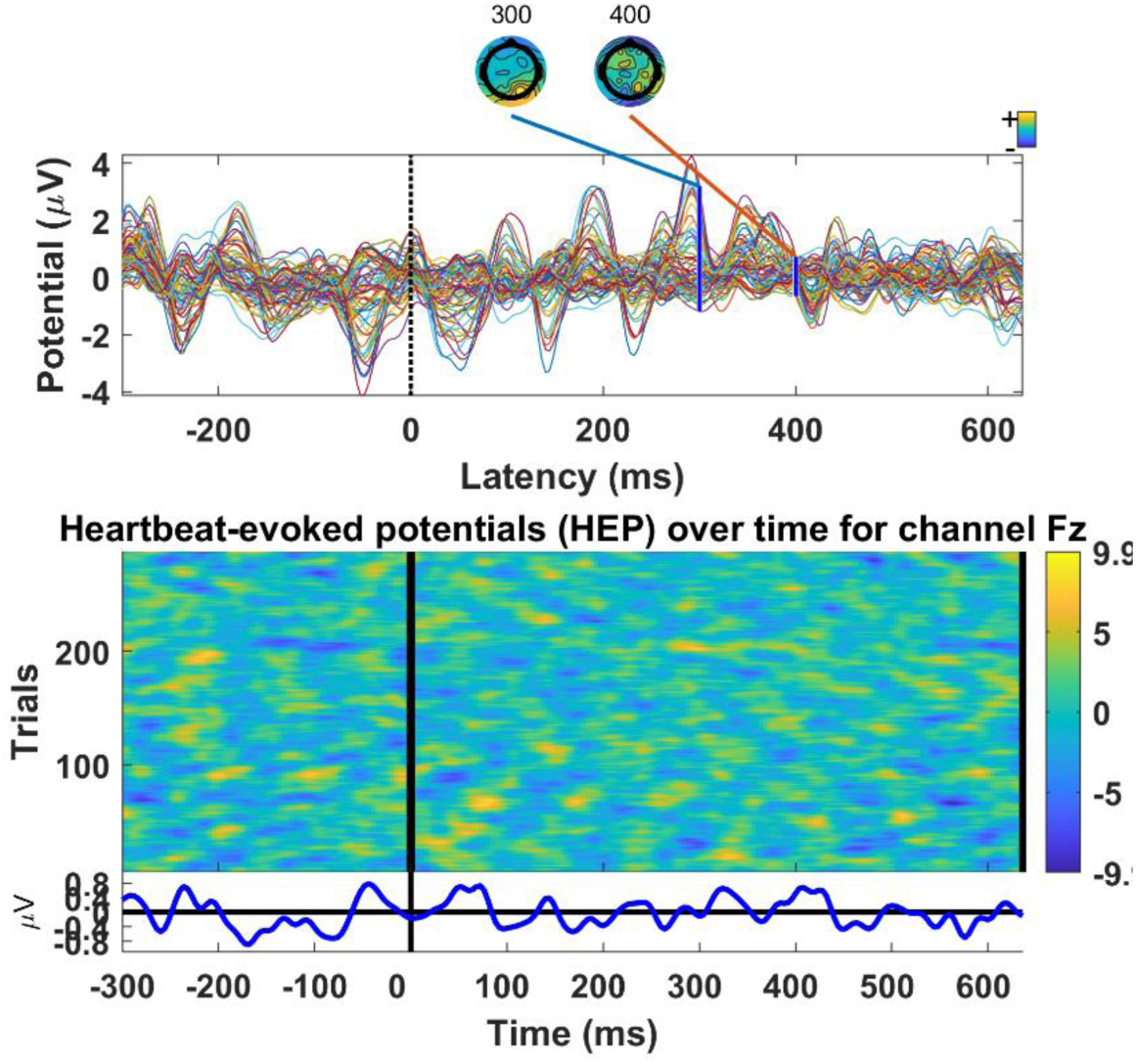
Grand average heartbeat-evoked potentials (HEP) obtained with PPG. **Top**: All electrodes are superimposed in the time domain, with scalp topographies showing amplitude distribution in the period of interest (200-500 ms after heartbeat). **Bottom**: HEP evolution over time (each “trial” corresponds to a pulse wave).

**Figure 18.**
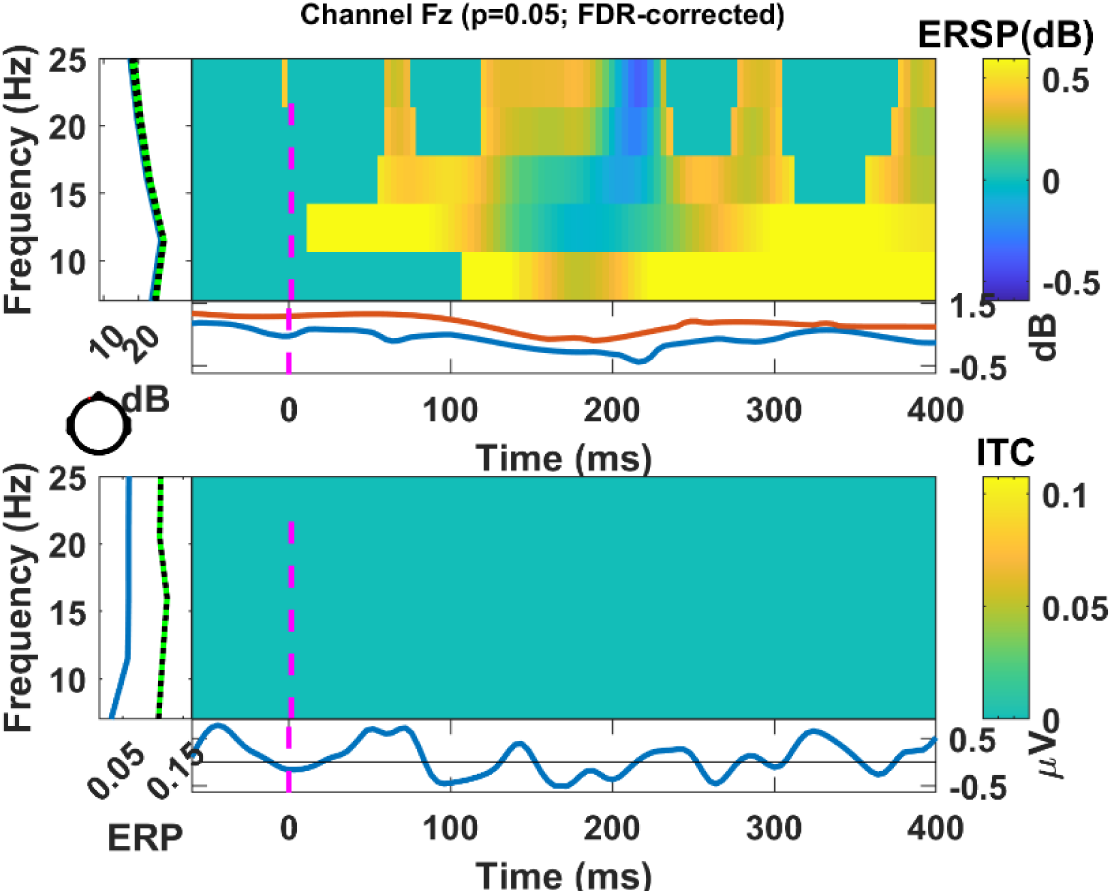
Heartbeat-evoked oscillations (HEO) obtained from PPG. **Top:** HEO for EEG channel Fz (frontocentral region) after permutation statistics (1000-iterations) and corrected for false discovery rate (FDR) at the 95% confidence level (p<0.05). **Bottom**: Inter-trial coherence (ITC) after FDR correction.

**Figure 19.**
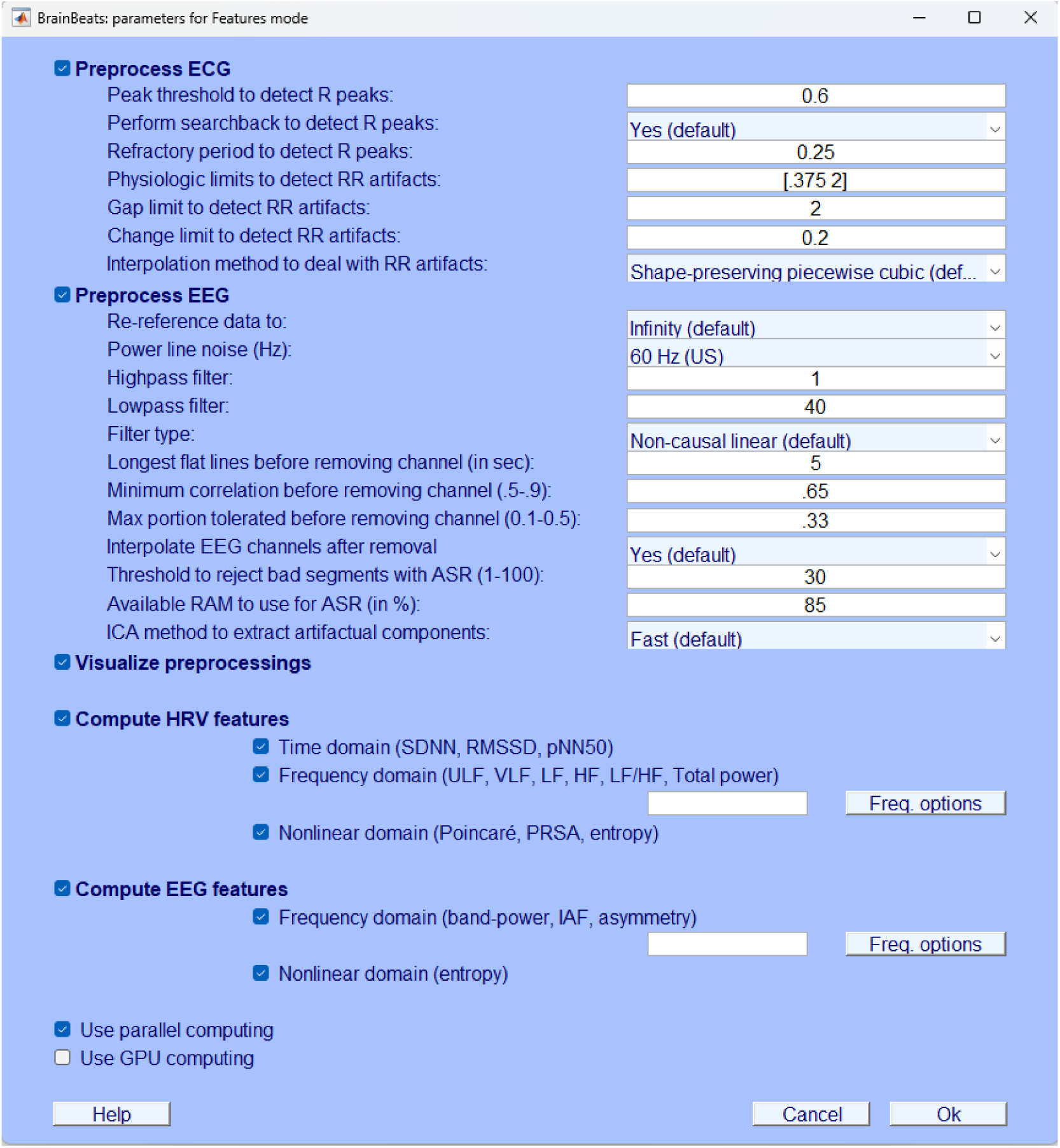
Selecting parameters for EEG & HRV feature extraction from ECG. This 2^nd^ graphical user interface (GUI) allows users to fine-tune parameters to 1) obtain the RR intervals from ECG signals, detect and interpolate the RR artifacts (if any) to obtain the NN intervals; 2) filter and re-reference EEG data, detect and interpolate abnormal EEG channels (if any), extract non-brain artifacts using ICA; 3) to visualize preprocessing steps; 4) compute heart-rate variability (HRV) features and in which domain (with additional parameters for the frequency domain by clicking on “freq. options”); 5) compute EEG features and in which domain (with additional parameters for the frequency domain by clicking on “freq. options”); 6) use parallel or GPU computing for faster computation time.

**Figure 20.**
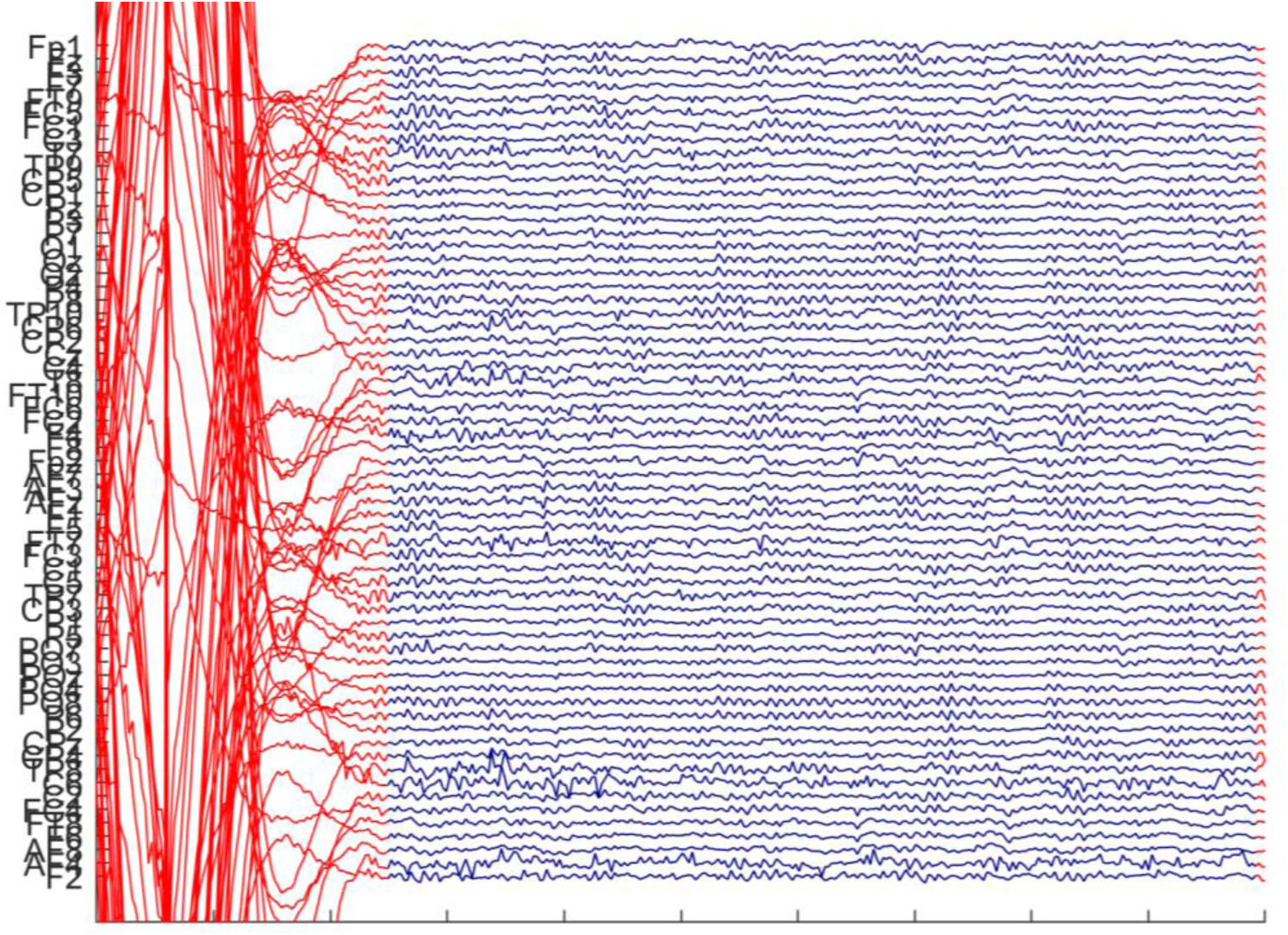
A large EEG artifact was detected and removed by the artifact subspace reconstruction (ASR) algorithm. Users can scroll through the whole file to inspect the segments removed by the algorithm.

**Figure 21.**
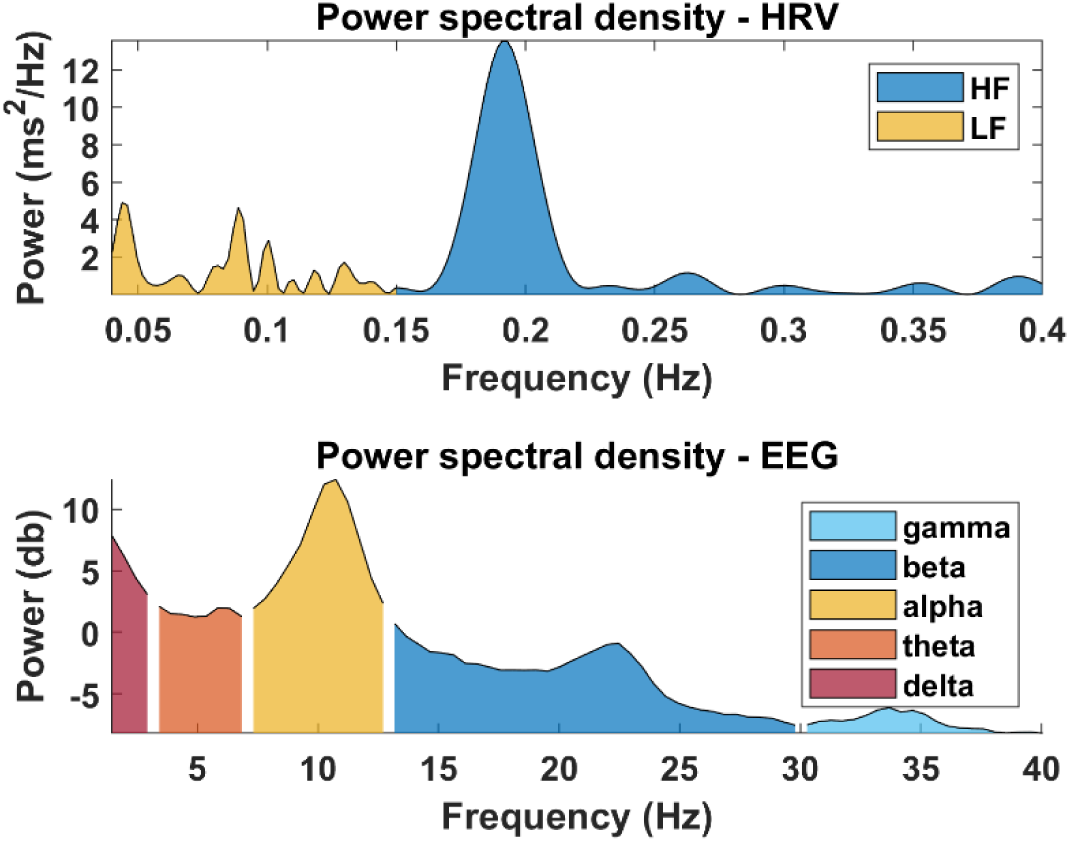
Power spectral density (PSD) extracted from NN intervals (ECG) and EEG signals. **Top**: Heart-rate variability (HRV) power in the low-frequency (LF; 0.04-0.15 Hz; in yellow) and high-frequency (HF; 0.15-0.40 Hz; in blue) bands, estimated using the normalized Lomb-Scargle periodogram from ECG signal. **Bottom**: PSD normalized to decibels (dB) calculated from the preprocessed EEG data, averaged across all channels for visualization.

**Figure 22.**
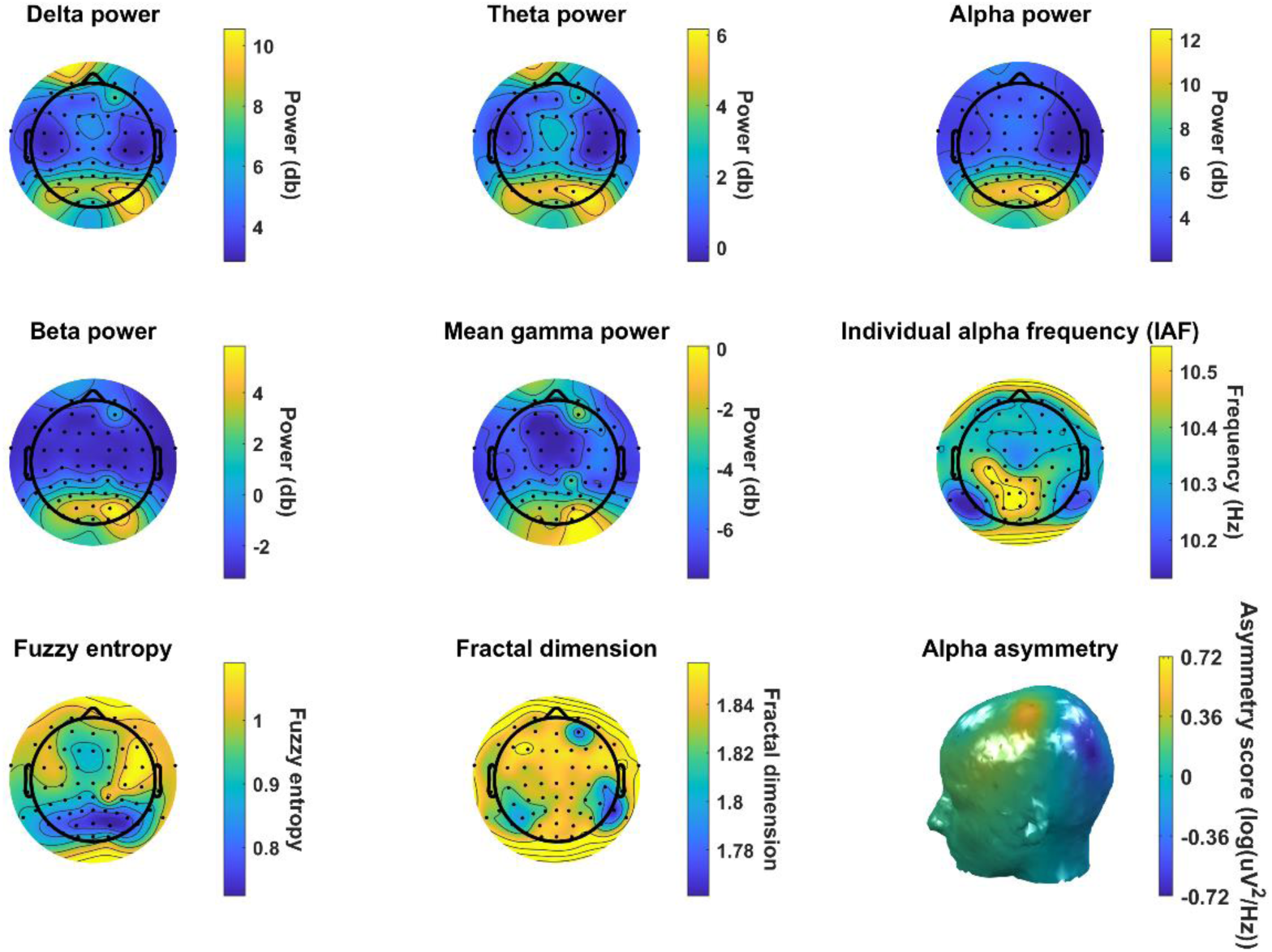
The main EEG features extracted by BrainBeats are illustrated by scalp topographies. The main EEG features include average spectral power in the delta (1-3 Hz), theta (3-7 Hz), alpha (8-13 Hz), beta (13-30 Hz), and gamma (30-40 Hz) frequency bands, the individual alpha frequency (IAF), fuzzy entropy, fractal dimension, and alpha asymmetry, in order. Notes: Greater fuzzy entropy values reflect higher complexity in terms of regularity, whereas the fractal dimension reflects greater complexity in terms of fractal characteristics. Alpha asymmetry was calculated on 16 symmetric pairs of electrodes. Positive values reflect greater left-than-right alpha power, typically associated with greater left-than-right inhibition of local cortical regions.

**Figure 23.**
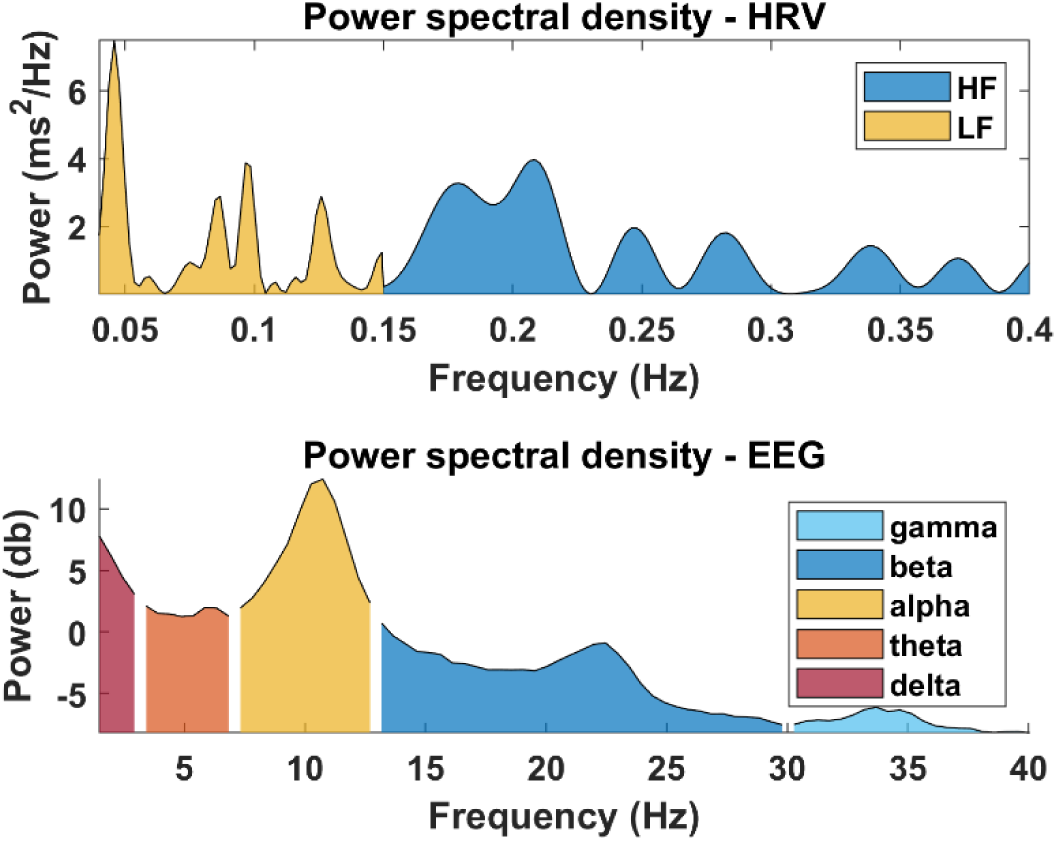
Power spectral density (PSD) extracted from NN intervals (PPG) and EEG signals. **Top**: Heart-rate variability (HRV) power in the low-frequency (LF; 0.04-0.15 Hz; in yellow) and high-frequency (HF; 0.15-0.40 Hz; in blue) bands, estimated using the normalized Lomb-Scargle periodogram from PPG signal.

**Figure 24.**
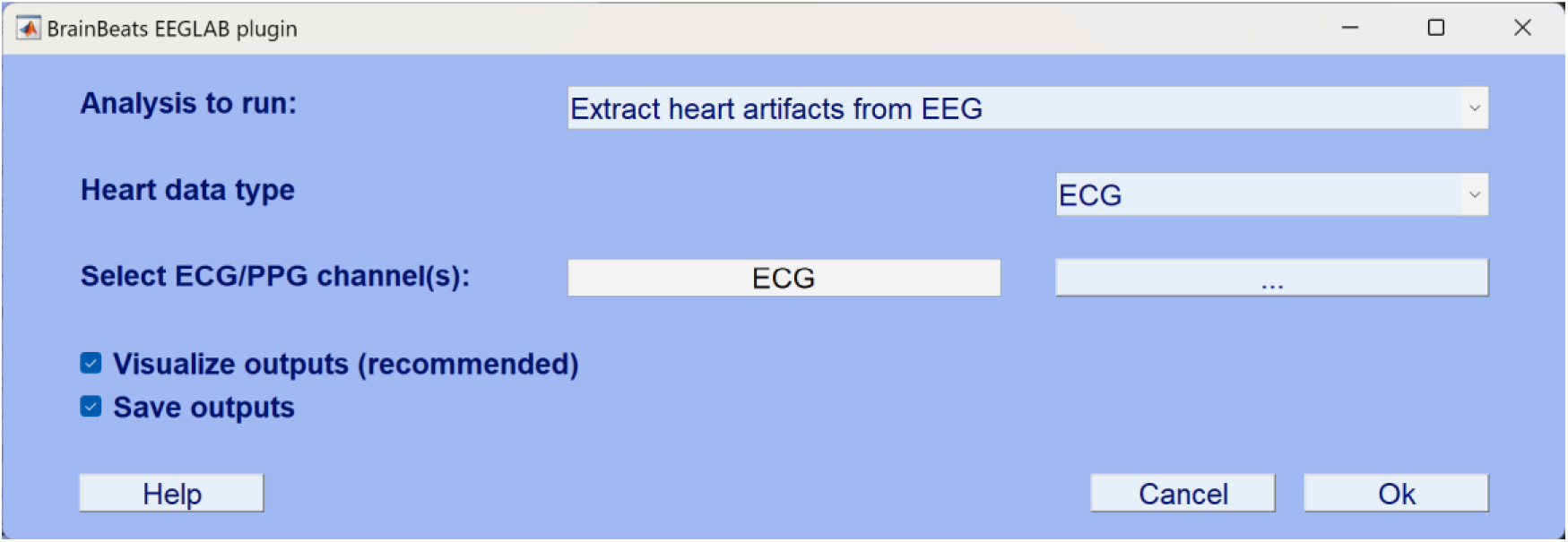
Selecting parameters for extracting heart artifacts from EEG.

**Figure 25.**
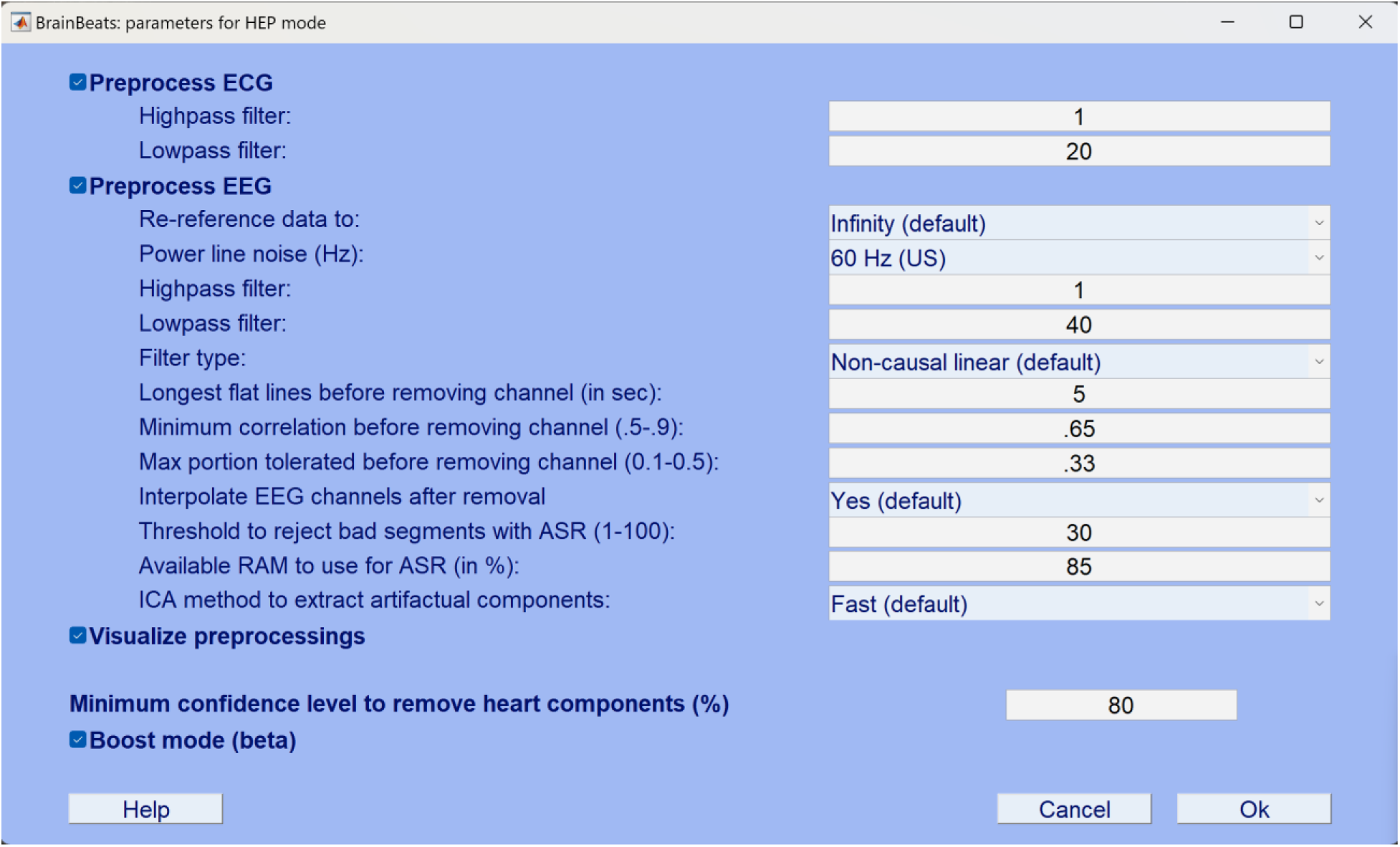
Fine-tuning parameters for extracting heart artifacts from EEG. In this 2^nd^ GUI, select preprocessing parameters prior to removing heart components from EEG signals. Data cleaning can dramatically affect the performance of this method. The confidence level to classify and remove heart artifacts is set to 80% by default. The “boost” mode can increase performance by smearing ECG signal across the EEG signals, but is turned off by default.

**Figure 26.**
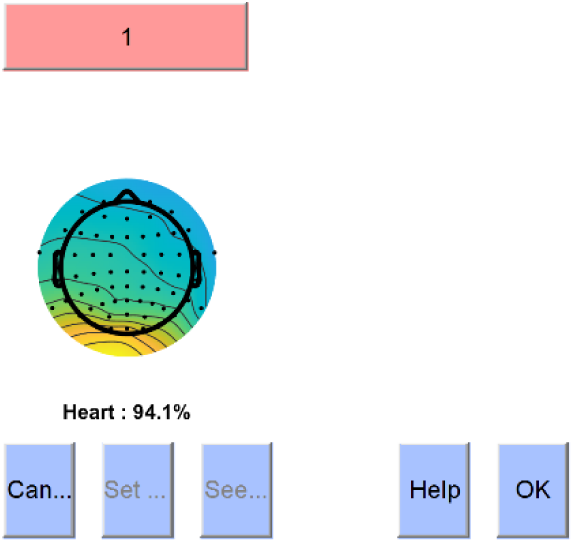
Visualization of the heart component detected and removed by BrainBeats.

**Figure 27.**
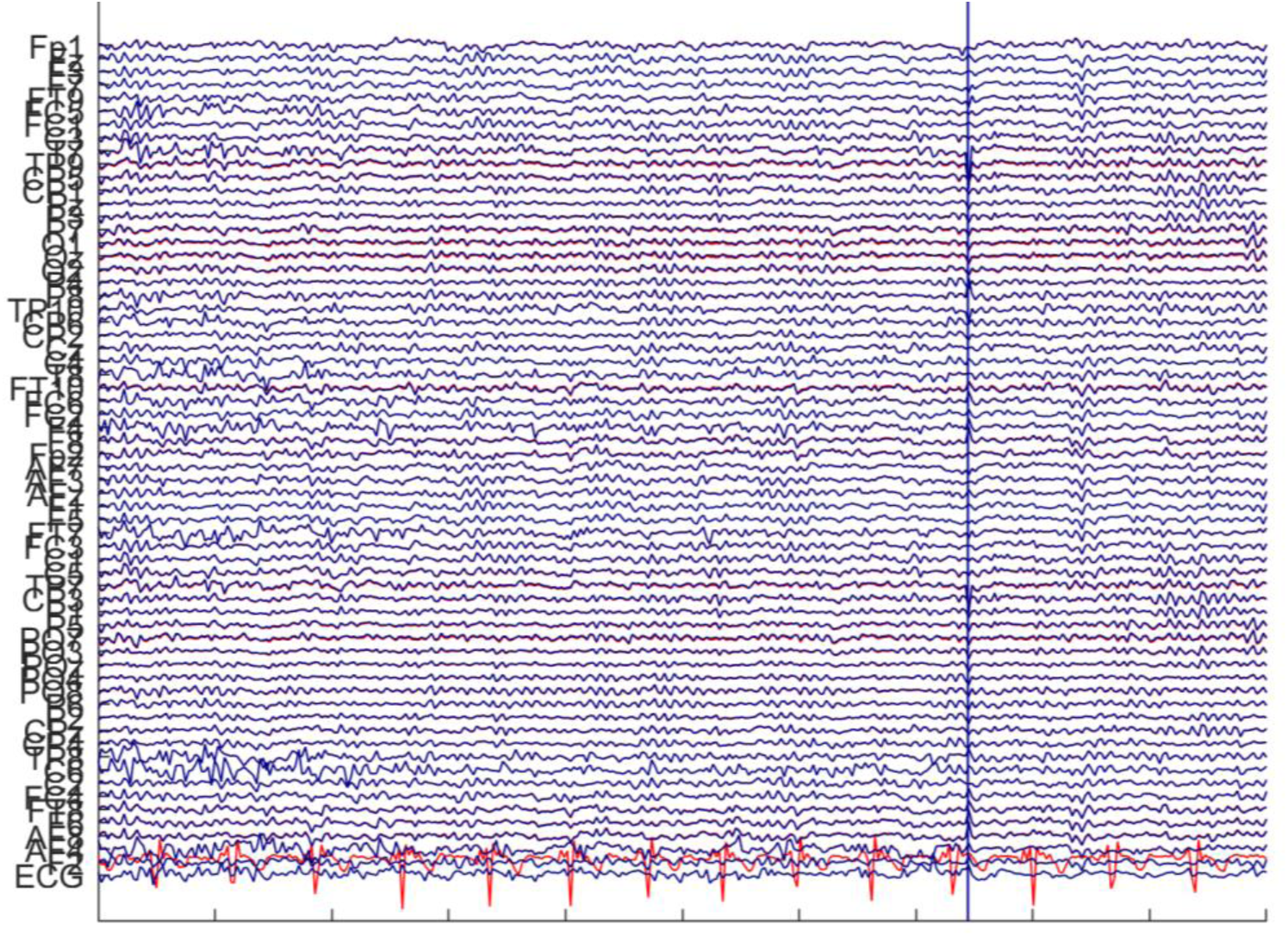
Visualization of the EEG signals after extracting heart artifacts. Visualize the final output of this method, i.e., the EEG signals (in blue) after extracting the ECG component (in red).

#### REPRESENTATIVE RESULTS

First, the BrainBeats plugin was used to preprocess EEG and ECG data, identify and remove artifacts, and analyze heartbeat-evoked potentials (HEP) and oscillations (HEO). BrainBeats successfully detected the RR intervals from the ECG signal and some RR artifacts (**Figure 4**). BrainBeats also reported in the command window that 11/305 (3.61%) of the heartbeats were flagged as artifacts and interpolated. The average signal quality index (SQI) of the RR intervals (before interpolation) has a value of 1, which is the highest value. A low signal quality corresponds to 20% or more of the RR series being flagged as an artifact or with an SQI<0.9. The TP9 EEG channel was removed and interpolated (**Figure 5**), several epochs containing artifacts (**Figure 8**), and two ocular components were flagged for removal with 99.9% confidence and 97.8% confidence, respectively (**Figure 9**). The output includes HEP waveforms for each EEG channel (**Figure 10**), grand average HEP (**Figure 11**), and grand average HEO (**Figure 12**). We observed significant heartbeat-evoked oscillations (HEO) in the alpha band (8-15 Hz) from 150 to 400 ms post-heartbeat at a frontocentral scalp site (**Figure 12, top**), consistent with previous findings ^17,67^). On the other hand, the inter-trial coherence (ITC) analysis suggested no significant phase-locking or resetting of the EEG phase with respect to the heartbeats (**Figure 12, bottom**).

Second, the BrainBeats plugin was used to do the same thing but with PPG. BrainBeats successfully detected the RR intervals from the ECG signal and some RR artifacts (**Figure 15**). BrainBeats reported in the command window that 15/309 (4.85%) of the heartbeats were flagged as artifacts and interpolated. The average signal quality index (SQI) of the RR intervals (before interpolation) has a value of 0.87. Signal quality is below 0.9 on average but still considered good because less than 20% of the RR series was flagged as an artifact or had an SQI<0.9. The outputs showed HEP waveforms for each EEG channel (**Figure 16**) and grand average HEP (**Figure 17**) based on the R waves detected from the PPG signal. We observed significant effects at a frontocentral scalp site in almost the whole time window following the R-wave (150-400 ms) and almost the whole frequency range (**Figure 18, top**). No ITC effect was observed (**Figure 18, bottom**). This is the first time HEP/HEO analysis has been performed from PPG signals, and future research is required to interpret these results.

Third, BrainBeats was used to extract EEG features and HRV features from the ECG signal. The process includes preprocessing the ECG and EEG signals, detecting and removing artifacts, and computing HRV and EEG features in various domains. We observed a peak in the HRV power spectral density (PSD) distribution at ∼0.19 Hz within the high-frequency (HF) band (**Figure 21, top**). For EEG, we observed a peak in the PSD distribution at ∼10.5 Hz within the alpha band **Figure 21, bottom**). The scalp topographies (**Figure 22**) indicate that the average power in the main frequency bands (and the highest peak alpha frequency values) are primarily localized in the posterior scalp areas. Furthermore, higher fuzzy entropy values (reflecting higher complexity in terms of regularity) are mostly localized in the frontal right and posterior scalp regions. In contrast, fractal dimension values (reflecting greater complexity in terms of fractal characteristics) do not show much variance across scalp regions. Finally, the alpha asymmetry plot (bottom right corner) shows greater left-than-right alpha power in the central parietal region and greater right-than-left alpha power in the posterior region. These differences in inter-hemispheric alpha power are generally interpreted in terms of local inhibition in the corresponding areas (i.e., more alpha power reflects greater cortical inhibition).

Fourth, BrainBeats was used to do the same, except that HRV features were extracted from the PPG signal. This time, we observe a peak at ∼0.04 Hz within the LF frequency band and a split peak around ∼0.19 Hz (**Figure 23, top**). Note that this distribution is slightly different than that obtained from the NN interval calculated from the ECG signal (**Figure 21, top**). This could be the result of lower signal quality in the PPG signal. EEG features are the same as in Figure 21.

Lastly, we used BrainBeats to extract heart artifacts from the EEG signals. A heart component was classified with 94.1% confidence using the “boost” mode (**Figure 26**), and extracted from the EEG signals (**Figure 27**).

## DISCUSSION

### Critical steps in the protocol

Critical steps are described in steps 1.1-1.4. Warnings and error messages are implemented at various places in the toolbox to help users understand why they may encounter issues (e.g., electrode locations not loaded in the EEG data, file length being too short for calculating a reliable measure of ultra-low frequency HRV, signal quality being too low for any reliable analysis, etc.). Each function is documented for advanced users, and the parameters can be easily fine-tuned (recommended parameters and typical ranges are documented in this manuscript and in the code). Users can access help regarding how to use certain functions or what normal ranges of parameters are, using the “help” command followed by the function’s name (e.g., type **help brainbeats_process** in the command window).

### Limitations of the method

Users must possess a dataset with EEG and cardiovascular (ECG or PPG) within the same file or know how to merge them independently. Data importation is currently not automated via BrainBeats because specific plugins need to be installed to account for the various data formats (e.g., .csv, .edf, .bdf, .vhdr, etc.).A future version will allow users to automatically load BIDS datasets with any number of files directly into BrainBeats, combine the EEG and cardiovascular signals available, and deal with potential time synchronicity issues (e.g., different sampling rates, align timestamps, etc.).

Entropy features are particularly promising for capturing complex, bidirectional interactions between cardiovascular, subcortical, and cortical systems that may be hidden in nonlinear feedback loop dynamics^27,73,74^. However, they are computationally heavy and can take a long time to compute from EEG signals, especially with high sampling rates. While some solutions are implemented (parallel computing, downsampling/decimation), future work will further reduce these computational costs.

While group-level statistics are available for HEP/HEO, they are not currently available for the features mode but will be available soon. Meanwhile, users may still use this method to extract features easily and reliably, perform statistics with any standard statistical software, or build feature-based ML models with their method of choice.

ECG directly assesses cardiac electrophysiology by capturing the heart’s electric fields. In contrast, PPG measures blood volume changes in microvascular beds, reflecting cardiac activity more indirectly through blood flow dynamics. Identifying the R-peak in ECG is straightforward due to its clear manifestation in the QRS complex, corresponding to the ventricular depolarization preceding the heart’s contraction. In contrast, the most prominent peak for the PPG signal is the systolic peak (or pulse wave peak), corresponding to the point of maximum blood volume in the arteries. It occurs slightly after the R-peak in the ECG. This delay is due to the time it takes for the pressure wave to travel from the heart to the peripheral site where the PPG signal is measured. Thus, the valley in the PPG waveform (marked as R-wave in BrainBeats), occurring between two systolic peaks, corresponds to the point of minimum blood volume and does not align with the R-peak in the ECG. Instead, it is closer to the T-wave in the ECG, which represents ventricular repolarization.^9^ This difference in signal characteristics results in timing disparities between ECG and PPG signals, which influence the temporal aspects of observed HEPs. Clinically, this divergence necessitates careful consideration in selecting the appropriate modality for HEP analysis, with ECG preferred for direct cardiac electrophysiological studies and PPG offering benefits in ease of use and patient comfort for long-term monitoring. While ECG and PPG can facilitate HEP analysis, their differing signal natures and physiological implications suggest that their inferences are not directly interchangeable. The choice between ECG and PPG for HEP analysis should be tailored to the specific objectives and needs of the study or clinical application. While this can be a limitation, it is also a strength because it means that ECG and PPG can provide two different types of information about the cardiovascular system, which can be complementary and provide new insights when combined. Furthermore, the temporal difference between R-peaks (from ECG) and R-waves (from PPG) could be corrected if stable over time, using a dataset containing simultaneous ECG-PPG signals as used for this tutorial.

PPG signals are smoother and lack distinct features relative to ECG, making them vulnerable to artifacts ^75^. While the algorithm used in this study was previously validated and performed well on the dataset used for this study, it might not perform as well on other types of PPG signals, especially those collected with wearable technologies.

For HEP/HEO analysis, epochs are defined using thresholds based on the distribution of the individuals’ interbeat intervals (IBI; ∼600-1000 ms). This leads to different epoch lengths across subjects and errors for group analysis. Furthermore, the short epoch length resulting from IBIs, relative to conventional EEG studies (stimuli are typically spaced in several seconds to allow the brain to return to baseline), leads to potential undesired edge effects or prevents users from examining slow frequencies. Time-frequency decomposition typically requires epochs to extend up to 3 cycles in the lowest frequency of interest beyond the window of interest. For HEO, the window of interest is 200-500 ms. Thus, one would require, for example, an additional 600 ms before and after the window (i.e., −400 to 900 ms) for examining frequencies as low as 5 Hz. If one wanted to examine frequencies as low as 1 Hz, an additional 3 s are needed before and after the window of interest. This is necessary for obtaining correct time and frequency resolution while avoiding edge effects.

### Significance of the method with respect to existing methods

Overall, BrainBeats provides state-of-the-art signal processing techniques for both EEG and cardiovascular signals, with fine-tuning capabilities via command line and graphical user interface (GUI).

The three methods can be performed using both a user-friendly graphical interface (GUI) and a command line (experts, allowing automation). Method 1 can be examined in the time (HEP), frequency (HEO), or time-frequency (HEO) domains, as well as at the channel or independent component levels. To our knowledge, HEP/HEO has never been performed using PPG signals and is now readily available.

Method 2 does not currently have a pre-existing alternative. Furthermore, users can also use BrainBeats to simply preprocess and extract EEG or HRV features individually with other datasets that do not contain both signals. For example, a user can preprocess ECG/PPG signals and extract HRV features to analyze HRV values only (and vice versa if a user wants to extract EEG features from an EEG dataset). This can be particularly useful for feature-based ML applications.

Method 3 allows quick and automated removal of heart artifacts from the EEG signals. While this is already possible in EEGLAB with the ICLabel plugin, it requires users to perform a series of steps and choice of parameters (e.g., highpass filtering the signals, running ICA, running ICLabel, tuning parameters, subtracting the heart components from the EEG signals, and removing the ECG channels) that can easily lead to errors (e.g., ghost ICs^55^). Furthermore, we introduce a “boost” method that increases the performance of this method using the cardiovascular signal (generally not included in these operations).

Additionally, the toolbox implements computing performance improvements to accelerate the estimation of EEG features (mainly multiscale entropy measures), including GPU and parallel computing. Note that these options are only as beneficial as the users’ hardware (i.e., graphic card and number of processors and threads).

### Future directions

The toolbox will continue to be modified and improved by the authors in the long term to implement field experts’ latest guidelines and recommendations and fix any errors that may arise.

Additional features and methods will be added to assess interactions between EEG and cardiovascular signals. For example, qEEG features such as theta/beta ratio (or similar spectral ratios that capture relevant clinical or cognitive information) can easily be added to the toolbox. New methods will include, for example, EEG-ECG direct and partial coherence, the time-resolved directional brain/heart interplay measurement^76^, or classification of HEP or feature data using machine learning (e.g., decision trees, random forest, naïve Bayes, SVM, KNN, long short-term memory networks, etc.)^17^.

For the best performance at detecting R-waves from noisy PPG signals collected with wearable technologies, future BrainBeats versions may provide alternative algorithms for these applications. Other promising algorithms include signal derivatives-based algorithms^77^, adaptive linear neuron artificial neural networks (used for ECG^78^), or ensemble empirical mode decomposition^79^.

For HEO analysis, to address the short epoch issue for time-frequency estimation (see limitations above), we plan to implement in future versions the reflection method, which mirrors the signal from the window of interest (i.e., backward version of the signal) before and after the window of interest to expand it. This provides smooth transitions and removes the undesired edge effects. The mirrored sections are then removed.

## ACKNOWLEDGMENTS

The Institute of Noetic Sciences supported this research. We thank the developers of the original open-source algorithms that were adapted to develop some of BrainBeats’ algorithms.

## REFERENCES

1. Bertalanffy, L. von. General System Theory: Foundations, Development, Applications. (G. Braziller, 1968).

2. Hodgkin, A. L. & Huxley, A. F. A quantitative description of membrane current and its application to conduction and excitation in nerve. J. Physiol. 117, 500–544 (1952).

3. Bean, B. P. Nitrendipine block of cardiac calcium channels: high-affinity binding to the inactivated state. Proc. Natl. Acad. Sci. U. S. A. 81, 6388–6392 (1984).

4. Fuchs, T. Ecology of the Brain: The Phenomenology and Biology of the Embodied Mind. (Oxford University Press, 2017).

5. Napadow, V. et al. Brain correlates of autonomic modulation: Combining heart rate variability with fMRI. NeuroImage 42, 169–177 (2008).

6. Chang, C., Cunningham, J. P. & Glover, G. H. Influence of heart rate on the BOLD signal: The cardiac response function. NeuroImage 44, 857–869 (2009).

7. Gianaros, P. J. & Sheu, L. K. A review of neuroimaging studies of stressor-evoked blood pressure reactivity: Emerging evidence for a brain-body pathway to coronary heart disease risk. NeuroImage 47, 922–936 (2009).

8. Harris, R. A history of electrocardiography: By George E. Burch, m.d. and Nicholas P. DePasquale, m.d. Year Book Medical Publishers, Inc., Chicago, Ill., 1964, pp. 309, $10. Am. J. Cardiol. 16, 769 (1965).

9. Allen, J. Photoplethysmography and its application in clinical physiological measurement. Physiol. Meas. 28, R1 (2007).

10. Cohen, M. X. Where Does EEG Come From and What Does It Mean? Trends Neurosci. 40, 208–218 (2017).

11. Cannard, C., Brandmeyer, T., Wahbeh, H. & Delorme, A. Self-health monitoring and wearable neurotechnologies. Handb. Clin. Neurol. 168, 207–232 (2020).

12. Gramann, K., Ferris, D. P., Gwin, J. & Makeig, S. Imaging natural cognition in action. Int. J. Psychophysiol. 91, 22–29 (2014).

13. Jungnickel, E., Gehrke, L., Klug, M. & Gramann, K. Chapter 10-MoBI—Mobile Brain/Body Imaging. in Neuroergonomics (eds. Ayaz, H. & Dehais, F.) 59–63 (Academic Press, 2019). doi:10.1016/B978-0-12-811926-6.00010-5.

14. Al, E. et al. Heart–brain interactions shape somatosensory perception and evoked potentials. Proc. Natl. Acad. Sci. 117, 10575–10584 (2020).

15. Banellis, L. & Cruse, D. Skipping a Beat: Heartbeat-Evoked Potentials Reflect Predictions during Interoceptive-Exteroceptive Integration. Cereb. Cortex Commun. 1, tgaa060 (2020).

16. Baranauskas, M., Grabauskaitė, A., Griškova-Bulanova, I., Lataitytė-Šimkevičienė, B. & Stanikūnas, R. Heartbeat evoked potentials (HEP) capture brain activity affecting subsequent heartbeat. Biomed. Signal Process. Control 68, 102731 (2021).

17. Candia-Rivera, D. et al. Neural Responses to Heartbeats Detect Residual Signs of Consciousness during Resting State in Postcomatose Patients. J. Neurosci. 41, 5251–5262 (2021).

18. Jiang, H. et al. Brain–Heart Interactions Underlying Traditional Tibetan Buddhist Meditation. Cereb. Cortex 30, 439–450 (2020).

19. Kumral, D. et al. Attenuation of the Heartbeat-Evoked Potential in Patients With Atrial Fibrillation. JACC Clin. Electrophysiol. 8, 1219–1230 (2022).

20. Thakor, N. V. & Tong, S. Advances in quantitative electroencephalogram analysis methods. Annu. Rev. Biomed. Eng. 6, 453–495 (2004).

21. Thayer, J. F., Åhs, F., Fredrikson, M., Sollers, J. J. & Wager, T. D. A meta-analysis of heart rate variability and neuroimaging studies: Implications for heart rate variability as a marker of stress and health. Neurosci. Biobehav. Rev. 36, 747–756 (2012).

22. Mather, M. & Thayer, J. F. How heart rate variability affects emotion regulation brain networks. Curr. Opin. Behav. Sci. 19, 98–104 (2018).

23. Kemp, A. H. & Quintana, D. S. The relationship between mental and physical health: Insights from the study of heart rate variability. Int. J. Psychophysiol. 89, 288–296 (2013).

24. Daneshi Kohan, M., Motie Nasrabadi, A., Shamsollahi, M. B. & Sharifi, A. EEG/PPG effective connectivity fusion for analyzing deception in interview. Signal Image Video Process. 14, 907–914 (2020).

25. Übeyli, E. D., Cvetkovic, D. & Cosic, I. Analysis of human PPG, ECG and EEG signals by eigenvector methods. Digit. Signal Process. 20, 956–963 (2010).

26. Zambrana-Vinaroz, D., Vicente-Samper, J. M., Manrique-Cordoba, J. & Sabater-Navarro, J. M. Wearable Epileptic Seizure Prediction System Based on Machine Learning Techniques Using ECG, PPG and EEG Signals. Sensors 22, 9372 (2022).

27. Shaffer, F. & Ginsberg, J. P. An Overview of Heart Rate Variability Metrics and Norms. Front. Public Health 5, 258 (2017).

28. Coan, J. A. & Allen, J. J. B. The state and trait nature of frontal EEG asymmetry in emotion. in The asymmetrical brain 565–615 (MIT Press, Cambridge, MA, US, 2003).

29. Hagemann, D., Hewig, J., Seifert, J., Naumann, E. & Bartussek, D. The latent state-trait structure of resting EEG asymmetry: replication and extension. Psychophysiology 42, 740–52 (2005).

30. Widge, A. S. et al. Electroencephalographic Biomarkers for Treatment Response Prediction in Major Depressive Illness: A Meta-Analysis. Am. J. Psychiatry 176, 44–56 (2019).

31. Olbrich, S. & Arns, M. EEG biomarkers in major depressive disorder: Discriminative power and prediction of treatment response. Int. Rev. Psychiatry 25, 604–618 (2013).

32. Kumar, Y., Dewal, M. L. & Anand, R. S. Epileptic seizures detection in EEG using DWT-based ApEn and artificial neural network. Signal Image Video Process. 8, 1323–1334 (2014).

33. Acharya, U. R. et al. Automated diagnosis of epileptic EEG using entropies. Biomed. Signal Process. Control 7, 401–408 (2012).

34. de Aguiar Neto, F. S. & Rosa, J. L. G. Depression biomarkers using non-invasive EEG: A review. Neurosci. Biobehav. Rev. 105, 83–93 (2019).

35. Cannard, C., Wahbeh, H. & Delorme, A. Electroencephalography correlates of well-being using a low-cost wearable system. Front. Hum. Neurosci. 15, 736 (2021).

36. Tarvainen, M. P., Niskanen, J.-P., Lipponen, J. A., Ranta-aho, P. O. & Karjalainen, P. A. Kubios HRV – Heart rate variability analysis software. Comput. Methods Programs Biomed. 113, 210–220 (2014).

37. Demski, A. J. & Soria, M. L. ecg-kit: a Matlab Toolbox for Cardiovascular Signal Processing. 4, e8 (2016).

38. Klug, M., et al. The BeMoBIL Pipeline for Automated Analyses of Multimodal Mobile Brain and Body Imaging Data. http://biorxiv.org/lookup/doi/10.1101/2022.09.29.510051 (2022) doi:10.1101/2022.09.29.510051.

39. Perakakis, P. HEPLAB. (2023).

40. Grosselin, F., Navarro-Sune, X., Raux, M., Similowski, T. & Chavez, M. CARE-rCortex: A Matlab toolbox for the analysis of CArdio-REspiratory-related activity in the Cortex. J. Neurosci. Methods 308, 309–316 (2018).

41. Luck, S. J. & Gaspelin, N. How to get statistically significant effects in any ERP experiment (and why you shouldn’t). Psychophysiology 54, 146–157 (2017).

42. Alday, P. M. How much baseline correction do we need in ERP research? Extended GLM model can replace baseline correction while lifting its limits. Psychophysiology 56, e13451 (2019).

43. Delorme, A. EEG is better left alone. Sci. Rep. 13, 2372 (2023).

44. Widmann, A., Schröger, E. & Maess, B. Digital filter design for electrophysiological data – a practical approach. J. Neurosci. Methods 250, 34–46 (2015).

45. Pham, T., Lau, Z. J., Chen, S. H. A. & Makowski, D. Heart Rate Variability in Psychology: A Review of HRV Indices and an Analysis Tutorial. Sensors 21, 3998 (2021).

46. Vest, A. N. et al. An open source benchmarked toolbox for cardiovascular waveform and interval analysis. Physiol. Meas. 39, 105004 (2018).

47. Smith, E. E., Reznik, S. J., Stewart, J. L. & Allen, J. J. B. Assessing and Conceptualizing Frontal EEG Asymmetry: An Updated Primer on Recording, Processing, Analyzing, and Interpreting Frontal Alpha Asymmetry. Int. J. Psychophysiol. Off. J. Int. Organ. Psychophysiol. 111, 98–114 (2017).

48. Dong, L. et al. MATLAB Toolboxes for Reference Electrode Standardization Technique (REST) of Scalp EEG. Front. Neurosci. 11, 601 (2017).

49. Candia-Rivera, D., Catrambone, V. & Valenza, G. The role of electroencephalography electrical reference in the assessment of functional brain–heart interplay: From methodology to user guidelines. J. Neurosci. Methods 360, 109269 (2021).

50. Mullen et al. Real-time Neuroimaging and Cognitive Monitoring Using Wearable Dry EEG. IEEE Trans. Biomed. Eng. Spec. Issue Wearable Technol. 62, 2553–67 (2015).

51. Chang, C.-Y., Hsu, S.-H., Pion-Tonachini, L. & Jung, T.-P. Evaluation of Artifact Subspace Reconstruction for Automatic EEG Artifact Removal. in 2018 40th Annual International Conference of the IEEE Engineering in Medicine and Biology Society (EMBC) 1242–1245 (2018). doi:10.1109/EMBC.2018.8512547.

52. Miyakoshi, M. Artifact subspace reconstruction: a candidate for a dream solution for EEG studies, sleep or awake. Sleep 46, zsad241 (2023).

53. Ablin, P., Cardoso, J.-F. & Gramfort, A. Faster independent component analysis by preconditioning with Hessian approximations. IEEE Trans. Signal Process. 66, 4040–4049 (2018).

54. Frank, G., Makeig, S. & Delorme, A. A Framework to Evaluate Independent Component Analysis applied to EEG signal: testing on the Picard algorithm. Preprint at http://arxiv.org/abs/2210.08409 (2022).

55. Kim, H. et al. ICA’s bug: How ghost ICs emerge from effective rank deficiency caused by EEG electrode interpolation and incorrect re-referencing. *Front*. Signal Process. 3, (2023).

56. Pion-Tonachini, L., Kreutz-Delgado, K. & Makeig, S. ICLabel: An automated electroencephalographic independent component classifier, dataset, and website. NeuroImage 198, 181–197 (2019).

57. Bigdely-Shamlo, N., Mullen, T., Kothe, C., Su, K.-M. & Robbins, K. A. The PREP pipeline: standardized preprocessing for large-scale EEG analysis. *Front*. Neuroinformatics 9, (2015).

58. Maris, E. & Oostenveld, R. Nonparametric statistical testing of EEG-and MEG-data. J Neurosci Methods 164, 177–90 (2007).

59. Pernet, C. R., Latinus, M., Nichols, T. E. & Rousselet, G. A. Cluster-based computational methods for mass univariate analyses of event-related brain potentials/fields: A simulation study. J Neurosci Methods (2015) doi:10.1016/j.jneumeth.2014.08.003.

60. Pernet, C. R., Chauveau, N., Gaspar, C. & Rousselet, G. A. LIMO EEG: A Toolbox for Hierarchical LInear MOdeling of ElectroEncephaloGraphic Data. Comput. Intell. Neurosci. 2011, (2011).

61 . Pernet, C. et al. Electroencephalography robust statistical linear modelling using a single weight per trial. *Aperture Neuro* 2022, (2022).

62. Pavlov, Y. G., Kasanov, D., Kosachenko, A. I., Kotyusov, A. I. & Busch, N. A. Pupillometry and electroencephalography in the digit span task. Sci. Data 9, 325 (2022).

63. Pavlov, Y. G., Kasanov, D., Kosachenko, A. I. & Kotyusov, A. I. EEG, pupillometry, ECG and photoplethysmography, and behavioral data in the digit span task and rest. Openneuro 10.18112/OPENNEURO.DS003838.V1.0.6 (2024).

64. Clifford, G. Signal processing methods for heart rate variability. (Oxford University, UK, 2002).

65. Pan, J. & Tompkins, W. J. A Real-Time QRS Detection Algorithm. IEEE Trans. Biomed. Eng. **BME****-**32, 230–236 (1985).

66. Maess, B., Schröger, E. & Widmann, A. Highpass filters and baseline correction in M/EEG analysis. Commentary on: “How inappropriate highpass filters can produce artefacts and incorrect conclusions in ERP studies of language and cognition”. J. Neurosci. Methods 266, 164–165 (2016).

67. Park, H.-D. & Blanke, O. Heartbeat-evoked cortical responses: Underlying mechanisms, functional roles, and methodological considerations. NeuroImage 197, 502–511 (2019).

68. Lomb, N. R. Least-squares frequency analysis of unequally spaced data. Astrophys. Space Sci. 39, 447–462 (1976).

69. Corcoran, A. W., Alday, P. M., Schlesewsky, M. & Bornkessel-Schlesewsky, I. Toward a reliable, automated method of individual alpha frequency (IAF) quantification. Psychophysiology 55, e13064 (2018).

70. Chen, W., Zhuang, J., Yu, W. & Wang, Z. Measuring complexity using FuzzyEn, ApEn, and SampEn. Med. Eng. Phys. 31, 61–68 (2009).

71. Cannard, C. & Delorme, A. An open-source EEGLAB plugin for computing entropy-based measures on MEEG signals. (2022) doi:10.31234/osf.io/xwmyk.

72. Lau, Z. J., Pham, T., Chen, S. H. A. & Makowski, D. Brain entropy, fractal dimensions and predictability: A review of complexity measures for EEG in healthy and neuropsychiatric populations. Eur. J. Neurosci. 56, 5047–5069 (2022).

73. Costa, M., Goldberger, A. L. & Peng, C.-K. Multiscale entropy analysis of biological signals. Phys. Rev. E Stat. Nonlin. Soft Matter Phys. 71, 021906 (2005).

74. Humeau-Heurtier, A. Multiscale Entropy Approaches and Their Applications. Entropy 22, 644 (2020).

75. Armañac-Julián, P. et al. Reliability of pulse photoplethysmography sensors: Coverage using different setups and body locations. *Front*. Electron. 3, (2022).

76. Catrambone, V., Greco, A., Vanello, N., Scilingo, E. P. & Valenza, G. Time-Resolved Directional Brain–Heart Interplay Measurement Through Synthetic Data Generation Models. Ann. Biomed. Eng. 47, 1479–1489 (2019).

77. Georgieva-Tsaneva, G., Gospodinova, E., Gospodinov, M. & Cheshmedzhiev, K. Portable Sensor System for Registration, Processing and Mathematical Analysis of PPG Signals. Appl. Sci. 10, 1051 (2020).

78. Kim, J.-H., Park, S.-E., Jeung, G.-W. & Kim, K.-S. Detection of R-Peaks in ECG Signal by Adaptive Linear Neuron (ADALINE) Artificial Neural Network. MATEC Web Conf. 54, 10001 (2016).

79. Lei, R., Ling, B. W.-K., Feng, P. & Chen, J. Estimation of Heart Rate and Respiratory Rate from PPG Signal Using Complementary Ensemble Empirical Mode Decomposition with both Independent Component Analysis and Non-Negative Matrix Factorization. Sensors 20, 3238 (2020).

